# Cannabinoids vs. whole metabolome: relevance of cannabinomics in analyzing *Cannabis* varieties

**DOI:** 10.1101/2021.06.07.447363

**Authors:** Pedro G. Vásquez-Ocmín, Guillaume Marti, Maxime Bonhomme, Fabienne Mathis, Sylvie Fournier, Stéphane Bertani, Alexandre Maciuk

## Abstract

*Cannabis sativa* has a long history of domestication both for its bioactive compounds and its fibers. This has produced hundreds of varieties, usually characterized in the literature by chemotypes, with Δ^9^-THC and CBD content as the main markers. However, chemotyping could also be done based on minor compounds (phytocannabinoids and others). In this work, a workflow, which we propose to name cannabinomics, combines mass spectrometry of the whole metabolome and statistical analysis to help differentiate *C. sativa* varieties and deciphering their characteristic markers. By applying this cannabinomics approach to the data obtained from 20 varieties of *C. sativa* (classically classified as chemotype I, II, or III), we compared the results with those obtained by a targeted quantification of 11 phytocannabinoids. Cannabinomics can be considered as a complementary tool for phenotyping and genotyping, allowing the identification of minor compounds playing a key role as markers of differentiation.

## 1. Introduction

*Cannabis sativa* L. (Cannabaceae) is an herbaceous annual plant cultivated since ancient times mainly in Central Asia and South-East Asia. It has been used as a source of fibers, foods, and its specific class of specialized metabolites, called cannabinoids, which are the basis for religious, recreational, and medicinal purposes [1]. Recent analysis suggests that *Cannabis* producing high levels of psychoactive compounds was smoked as part of ritual activities in western China at least 2500 years ago [2]. So far, over 600 constituents have been reported in *Cannabis* plants, including ≈ 200 terpenes (mono-, di-, sesqui-and triterpenes), ≈ 25 flavonoids, ≈ 150 cannabinoids, and other compounds like stilbenes, lignans, phytosterols, alkaloids, and amides [3,4]. Cannabinoids from *C. sativa*, localized in glandular trichomes, are also called phytocannabinoids to distinguish them from the mammalian endogenous (endocannabinoids) or synthetic compounds able to bind to cannabinoid receptors. These terpenophenolic compounds (i.e., isoprenylated polyketides) are considered as the active constituents of *Cannabis* plants and may contain 19 to 23 carbons depending on the length of the alkyl side chain [5]. They originate from the convergence of two distinct biosynthetic pathways: the polyketide pathway, giving rise to olivetolic acid (OLA) or its C1, C5 or C7 analogs, and the plastidial methyl-D-erythritol 4-phosphate (MEP) pathway, leading to the prenylation of OLA with geranyl diphosphate (GPP) that undergoes oxidative cyclization reactions [6,7]. Δ^9^-tetrahydrocannabinolic acid (Δ^9^-THCA), cannabidiolic acid (CBDA), cannabichromenic acid (CBCA), and their common precursor cannabigerolic acid (CBGA) are the major phytocannabinoids found natively in most of the varieties. These acidic cannabinoids are rather unstable and may spontaneously lose the carboxyl group with exposure to light, air or heat. Their decarboxylated analogs like cannabidiol (CBD) or Δ^9^-tetrahydrocannabinol (Δ^9^-THC) show a different, and often higher, bioactivity [8,9]. Many phytocannabinoids were characterized between the 1960s and 1980s [10–12], although new cannabinoids, sometimes showing cannabimimetic action, are continuously identified [5,13–15]. *Cannabis* chemotypes (also called phenotypes or chemovars) are commonly split into three groups based on their Δ^9^-THC/CBD ratio: Δ^9^-THC/CBD >> 1 (chemotype I, “drug-type”), Δ^9^-THC/CBD ≥ 1 (chemotype II, “intermediate-type”), Δ^9^-THC/CBD << 1 (chemotype III, “fiber-type” or “hemp”) [16–18]. Additional chemotyping can also be based on the terpenoids, with at least five chemotypes described [19–21]. Hemp usually contains significant amounts of non-intoxicating cannabinoids, *e.g.* cannabidiolic acid (CBDA) or cannabigerolic acid (CBGA). Although environmental factors play a key role in determining the amount of phytocannabinoids present in the plant at different growth stages, the Δ^9^-THC/CBD ratio is under genetic control and is quite stable for one given chemotype [22].

To date, medicinal use of *Cannabis* is authorized in more than 50 countries, while its recreational use is authorized in around 20 countries. In December 2020, The UNODC Commission on Narcotic Drugs removed *Cannabis* from Schedule IV, which list drugs being particularly liable to abuse and to produce ill effects, to incorporate it into Schedule I, which allows its therapeutic use [23]. *Cannabis* and isolated phytocannabinoids have shown beneficial effects for chronic pain, multiple sclerosis, neurodegenerative disorders, treatment-resistant epilepsy, nausea and vomiting associated with chemotherapy diseases, cancer, cardiovascular disorders, metabolic disorders and more [24–28]. Besides its current move towards medicinal *Cannabis* through an experimental status for 2 years (2021-2023), France has been for decades a leading producer of hemp in Europe. The commercial hemp seed producer (HEMP-it, formerly known as *Fédération Nationale des Producteurs de Chanvre*) holds a large number of *Cannabis* fiber-type varieties [29–33].

The scientific literature on *Cannabis* has grown at a staggering pace, with 10 articles published per day. Yet, a majority of them pertain to Δ^9^-THC and CBD and to far less extent the varin type Δ^9^-THCV, leaving the remaining metabolites somewhat in the shadows. Chemical differentiation of *Cannabis* chemotypes is currently done by using 2 to 15 major cannabinoids as principal markers. Meanwhile, comprehensive metabolomic approaches allowing the detection and annotation of numerous metabolites are currently becoming mainstream in plant science and have become a key to decipher their biological roles. Techniques used for detection are hyphenated liquid or gas chromatography-mass spectrometry (LC- or GC-MS) or nuclear magnetic resonance spectroscopy (NMR), both techniques being assisted by bioinformatics and statistical analysis. We applied a LC-MS-based metabolomics workflow on medium polarity extracts of *C. sativa* leaves and flowers to classify chemotypes as well as map and annotate metabolites of the phytocannabinoids family and other phytochemical classes. In a collection of 20 genetically diverse chemotypes, we compared the ability of this approach to decipher chemotypes markers with an approach based only on major cannabinoids quantification.

## 2. Material and methods

All solvents used for chromatography were LC-MS grade (HiPerSolv Chromanorm, VWR, France, Fontenay-Sous-Bois). Milli-Q RG system (Millipore, France) was used to produce high purity water of 18.2 MΩ.cm resistivity. Formic acid was >98% for LC-MS (Fluka, Buchs, Switzerland). Phytocannabinoids standards (CBDA, CBGA, CBCA, Δ^9^-THCA, CBD, CBG, CBC, Δ^9^-THC, Δ^8^-THC, THCV, and CBN) were purchased from Sigma-Aldrich (St. Louis, MO, USA) and immediately stored at −30 °C.

### 2.1. Plant material

Thirteen fiber-type (chemotype III) and three intermediate-type (chemotype II) *Cannabis sativa* varieties were obtained from the HEMPT-it ADN company (Beaufort-en-Anjou, France) and four drug-type varieties (chemotype I) were obtained from CBDIS company (Paris, France) (see the list and description of varieties in **supp info 1**). Chemotype III and II samples were dried at 40°C, while chemotype I was dried at 80°C; all samples were finely ground, then stored at room temperature.

### 2.2. Metabolome analysis

For each sample, 100 mg were extracted in triplicate using 5 mL of acetonitrile/methanol (8:2 v/v) under sonication for 30 minutes. Extracts were then centrifuged at 4000 g for 15 minutes. Supernatants were filtered using PTFE membrane syringe filter, (50 mm × 0.2 μm) and stored in vials at −30 °C. A quality control (QC) sample was created by pooling aliquots from each extract. Extracts were diluted 10 times and standards prepared at 10 μg/mL in the same solvent. Ultra-High Performance Liquid Chromatography – High-Resolution MS (UHPLC−HRMS) analyses were performed on a Q Exactive Plus orbitrap mass spectrometer, equipped with a heated electrospray probe (HESI II) coupled to a U-HPLC Ultimate 3000 RSLC system (Thermo Fisher Scientific, France). Separation was done on a Luna Omega Polar C18 column (150 mm × 2.1 mm i.d., 1.6 μm, Phenomenex, Sartrouville, France) equipped with a guard column. The mobile phase gradient used water containing 0.05 % formic acid (A) and acetonitrile containing 0.05% of formic acid (B). A gradient method at a constant flow rate of 0.3 mL/min was applied under the following conditions: 60 to 100 % B in 20 min, 100 % B over 2 min, equilibration over 4 min. Column oven temperature was set to 30°C, autosampler temperature was set to 5°C, and the injection volume was fixed to 2μL. Mass detection was performed in positive ionization (PI) mode at resolution 30,000 [fullwidth at half-maximum (fwhm) at 400m/z] for MS1 and resolution 17,500 for MS2 with automatic gain control (AGC) target of 1×10^6^ for full scan MS1 and 1×10^5^ for MS2. Ionization spray voltage was set to 3.5 kV for PI, and the capillary temperature was set to 256 °C. The mass scanning range was *m/z* 100−1500. Each full MS scan was followed by a data-dependent acquisition of MS/MS spectra for the six most intense ions using stepped normalized collision energy of 20, 40, and 60 eV.

### 2.3. Data mining process

LC-MS data were processed following the MSCleanR workflow [34]. Briefly, samples were processed with MS-DIAL version 4.38 [35]. MS1 and MS2 tolerances were set to 0.01 and 0.05 Da, respectively, in centroid mode for each dataset. Peaks were aligned on QC reference with a RT tolerance of 0.2 min, a mass tolerance of 0.015 Da, and minimum peak height detection at 4.5 × 10^6^. MS-DIAL data were deconvoluted with MS-CleanR by selecting all filters with a minimum blank ratio set to 0.8, and a maximum relative standard deviation (RSD) set to 40. The maximum mass difference for feature relationships detection was set to 0.005 Da, and the maximum RT difference was set to 0.025 min. The Pearson correlation links were applied for correlation ≥ 0.8 and *p*-value significance threshold = 0.05. Two peaks were kept in each cluster for further request and the kept features were annotated with MS-FINDER version 3.50 [36]. The MS1 and MS2 tolerances were set to 5 and 15 ppm, respectively. Formula finder was exclusively processed with C, H, O, N, atoms. For compounds annotation, several sets of data were used: 1) experimental LC-MS/MS data of 11 phytocannabinoid standards, analyzed on the same instrument and with the same method, were used as references using MSDial based on accurate and exact mass, fragmentation, retention time; 2) compounds identified in the *Cannabis* genus and the Cannabaceae family, generated from literature data (Dictionary of Natural Products, DNP on DVD v. 28.2, CRC press) were mined by MS Finder based on exact mass and *in-silico* fragmentation; 3) a generic database encompassing exact mass and MS/MS data of compounds from natural products databases included in MS-FINDER (KNApSAcK, PlantCyc, ChEBI, NANPDB, UNPD, and COCONUT). Statistical analysis and mass spectra similarity network (molecular network) was carried out using the open-source software MetGem [37] on final .msp and .csv files obtained with MSCleanR. Values used were MS2 *m/z* tolerance = 0.05 Da, minimum matched peaks = 4 and minimal cosine score value = 0.7. Visualization of the network was performed on Cytoscape version 3.8.2 [38].

### 2.4. Genotyping

Fifteen cultivars from chemotypes II and III (excepted Epsilon variety) were studied with 24 plants per cultivar to perform genetic analyses. DNA extraction was performed with the Macherey nagel NucleoSpin Plant II kit from foliar discs taken from young plant leaves. Sixteen SSR markers (see **supp info 2**) were chosen for their polymorphism and repeatability [39,40]. The PCR MIX contains 10 ng of genomic DNA, 250 μM dNTP, 0.2 μM of primer, 2.0 mM of MgCl2, 2.0 μg.μl^−1^ of BSA, 10 mM of PCR buffer, and 0.3 units Taq Gold polymerase (Perkin Elmer). PCRs consisted of a 5 min preheat at 95°C followed by two rounds of cycles. Touchdown PCR consisted of 10 cycles of 95°C denaturation for 15 s, 63°C touchdown to 53°C annealing for 30 s, and 72°C extension for 30 s. This was followed by 19 cycles of 95°C denaturation for 15 s, 52°C annealing for 30 s, 72°C extension for 1 min, and one 72°C final extension for 10 min.

### 2.5 Quantification of main phytocannabinoids

Mixes of 11 phytocannabinoids at six final concentrations (10000, 1000, 100, 10, 1, 0.1, and 0.001 ng/mL) were performed to obtain calibration curves for quantitative analysis. All extracts were profiled using the LC-MS method described above and were processed using MZmine 2.53 [41] Briefly, after mass detection, the chromatogram was built and deconvolutioned using ADAP chromato-builder and wavelets (ADAP) algorithms [42]; a grouping of isotope patterns (peak grouper algorithm) and a unique pick list aligned was created; finally, a gap-filling (peak finder algorithm) was performed. A final list of the peak height for each standard at each concentration was carried out and a calibration curve was created for each standard. Validation parameters were determined following the International Conference on Harmonization (ICH) Guidelines [43], and the following characteristics were evaluated: linearity, limit of detection (LOD), and limit of quantification (LOQ).

### 2.6 Statistical and genetic analyses

The final annotated metabolome dataset generated by the MS-CleanR workflow was uploaded on the MetaboAnalyst 5.0 online platform [44]. The data were normalized by sum and scaled by unit variance before statistical analysis. First, principal component analysis (PCA) was applied as an exploratory data analysis to provide an overview of LC-MS fingerprints. Then, a hierarchical cluster analysis (HCA) was performed to obtain a dendrogram of varieties using the metabolome dataset and the phytocannabinoids quantification dataset. Briefly, for each variable the average value across replicates was calculated, then distance matrices were obtained between the 20 varieties using either 1-*r* (the Pearson correlation coefficient) for the metabolome dataset or the Euclidean distance for the phytocannabinoids quantification dataset, followed by HCA on each matrix using Ward’s minimum variance criterion. On the clusters obtained, a partial least squares discriminant analysis (PLS-DA) was conducted using clusters as Y value and the top 50 features were plotted on a heatmap using ANOVA. Additionally, to look for specific biomarkers of each variety the same statistical approach was applied within each cluster.

Genetic differentiation among fifteen *Cannabis* varieties based on 16 microsatellite markers was measured by *F_ST_* using the R package *adegenet*, and visualized by HCA using Ward’s minimum variance criterion. To evaluate the degree of genetic structuring of cannabis varieties without *a priori*, we used the software STRUCTURE v2.3.4 [45], which implements a Bayesian algorithm to identify *K* user-defined clusters of genetically homogeneous individuals. We analyzed the data with *K* = 2 to 20. The analysis produced for each individual a proportion of membership to each cluster. We used both the ‘‘admixture’’ and ‘‘no admixture’’ models. Allelic frequencies were set to correlate among populations. All analyses were replicated 10 times to ensure proper convergence of the MCMC with a burn-in of 100 000 steps and a MCMC length of 1 000 000 after burn-in. Posterior inference of the optimal number of cluster was performed using the *ad hoc* statistic Δ*K* based on the likelihood [46].

## 3. Results and Discussion

### 3.1. Cannabinomics reveals major discriminant compounds between *Cannabis* varieties

Identification of phytocannabinoids in *Cannabis* plants is difficult for several reasons: only a few of them are commercially available, those that are available ones are expensive to purchase, access to some standards is restricted by national regulations, and the carboxylic analogs are of highly labile nature. Therefore, most studies encompass only a handful of standards. An increasing number of studies use mass spectral libraries (*i.e.* NIST20 library) [47] or in-house databases [48] for the annotation of the *Cannabis* metabolome [47], in a metabolomic approach variously referred to as Cannabinomics, Cannabinoidomics, or Phytocannabinomics [49–51]. We used several mass spectral libraries and three levels of annotations (see above in **2.3**), allowing in most cases an annotation of level 2 according to the metabolomic standard initiative [52]. Application of the MSCleanR workflow to the entire LC-MS dataset in positive mode resulted in 396 compounds. Annotation prioritization was done by ranking from highest to lowest confidence as follows: 1) standards, 2) genus and family DB, and 3) generic DB. Among the compounds detected, 193 had a match in the *Cannabis* genus (48.7%), 41 in the Cannabaceae family (10.4%), 84 in the generic database (21.2%) while 78 remained unannotated (19.7%) (**Table in Supp info 3**). Extraction with ACN/MeOH 8:2 v/v is expected to yield the medium and low polarity parts of the metabolome, including phytocannabinoids and non-polar compounds. The set of extracts allowed identification or annotation of 105 phytocannabinoids under acidic or neutral form (26.5% of compounds annotated). Other metabolites annotated mostly included flavonoids and terpenoids (mono-, sesqui-, triterpenoids, prenol lipids) (**Figure 1**). Unknown compounds distributed evenly along the chromatogram and their masses spaned in the *m/z* 300-1200 range. Structural similarity between compounds, assessed *via* fragmentation pattern similarity, is reflected by the molecular network (MN) **(Figure 1 and supp info 4)**. Phytocannabinoid standards dispatched into 4 clusters, following pathways of fragmentations consistent with the previous report of Berman *et al.* [53]. One cluster gathers neutral forms of THC and CBD, with several compounds being annotated as isomers of THC (Δ^8^-, Δ^7^-, Δ^4^-, Δ^4^-^8^-, Δ^4^-, Δ^4^-THC), as well as C3-derivatives (varin-type). Another cluster gathers acidic derivatives of THC, CBD and CBC. Both neutral and acidic derivatives of CBG-type cannabinoids are grouped together in another cluster. Four compounds are grouped with CBN in a cluster that contains degradation products, annotated as cannabinodiol isomers and cannabinol hydroxylated on the side chain. It is noteworthy that CBE derivatives (degradation products of CBD) are clustered with CBG derivatives. Cannabicyclolic acid (CBL) derivatives, which are degradation products of cannabichromene (CBC), are clustered in a group containing 14 compounds.

**Figure 1.**
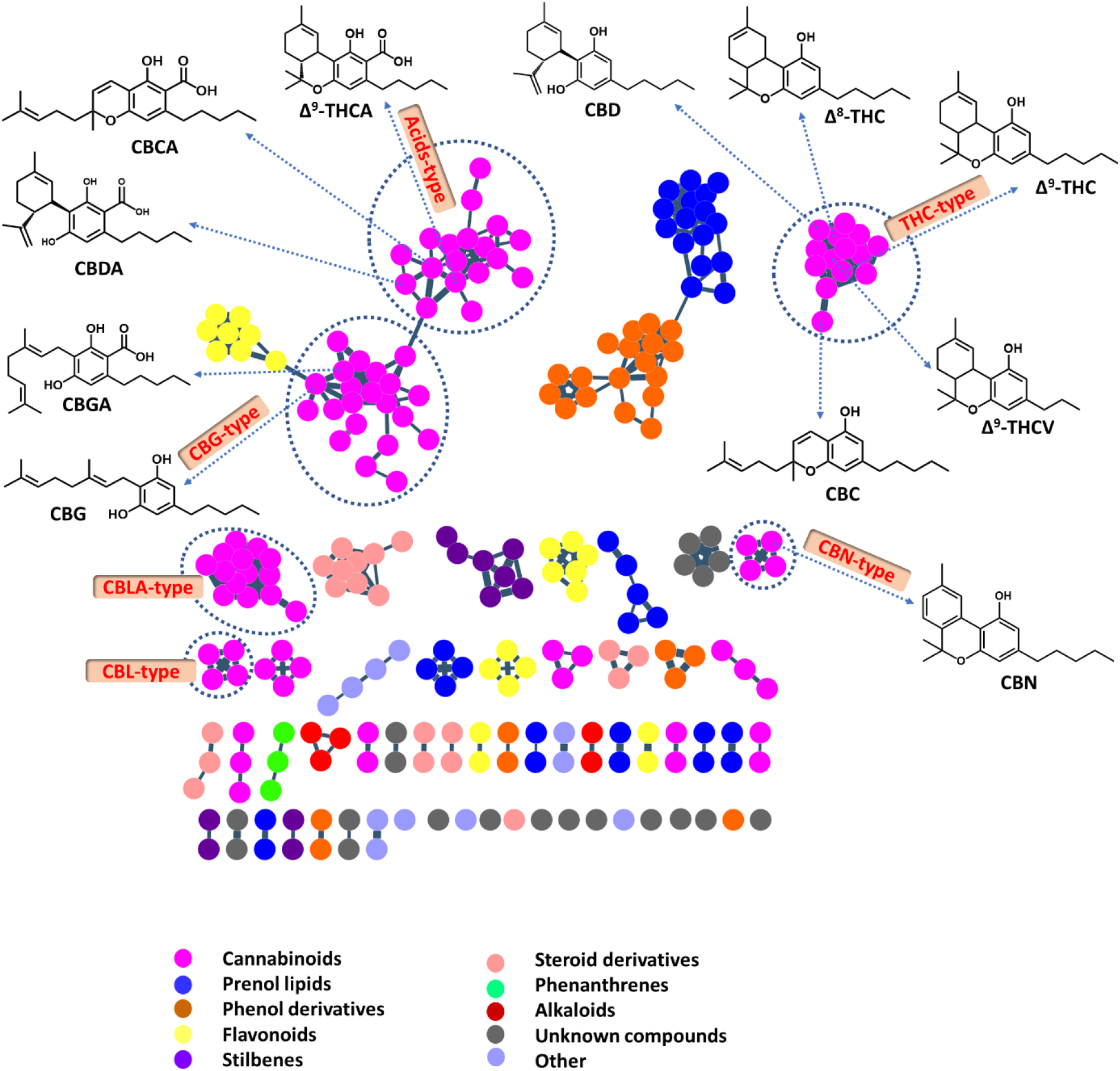
Mass spectral similarity network (molecular network) of annotated compounds based on MetGem software and visualized by Cystoscape. Colors show the major class of compounds in the metabolome for all *Cannabis* varieties, based on Classyfire attribution [54] and MetGem clustering.

Currently, more than 600 *Cannabis* varieties are commercially available whose genetics, for many of them, is only partially known [49]. In this work, we characterize twenty varieties of *Cannabis* (sixteen hemp varieties and four recreative varieties) using metabolomic tools. In order to visualize relationships between varieties based on their genotypes and phenotypes, hierarchical cluster analyses (HCA) were performed on each set of data (**Figure 2-A to C**). HCA based on genetic differentiation among varieties (*F_ST_*) suggests a clustering into two main genetic groups (showed in red and green, **Fig 2-A**) as confirmed by the STRUCTURE analyses (**supp info 5**). Most of the varieties grouped together (in green) belong to a pool of varieties developed by Hemp-It by cross-breeding and massal selection, still showing a substantial amount of genetic diversity. Clustering based on LCMS profiles on a wider panel of varieties (**Fig 2-B** and **2-C**) yields a very similar clustering for both global metabolome data and quantified main cannabinoids data. These first analyses validate the long-standing literature classifying *C. sativa* varieties based on the THC/CBD ratio, leading to 3 chemotypes: hemp-type (chemotype III), drug-type (chemotype I), and intermediate type (chemotype II) [16–18]. Among the varieties we studied, chemotype III encompasses **cluster 3** (CBD-majority varieties) and **1** (CBG-majority varieties), whereas chemotype II is illustrated by varieties used for seed production (**cluster 2**). Drug-type varieties (chemotype I) are clearly clustered out (**cluster 4**). Samples of chemotype III were developed by HEMP-it (French hemp seed producer) in a dedicated work based on GC and TLC analyses to obtain non-monoclonal varieties with THC content below 0.2 % [33]. While selection criterion produced a homogeneous chemotype III, minor metabolites may nevertheless be used to distinguish sub-varieties, as we show below.

**Figure 2.**
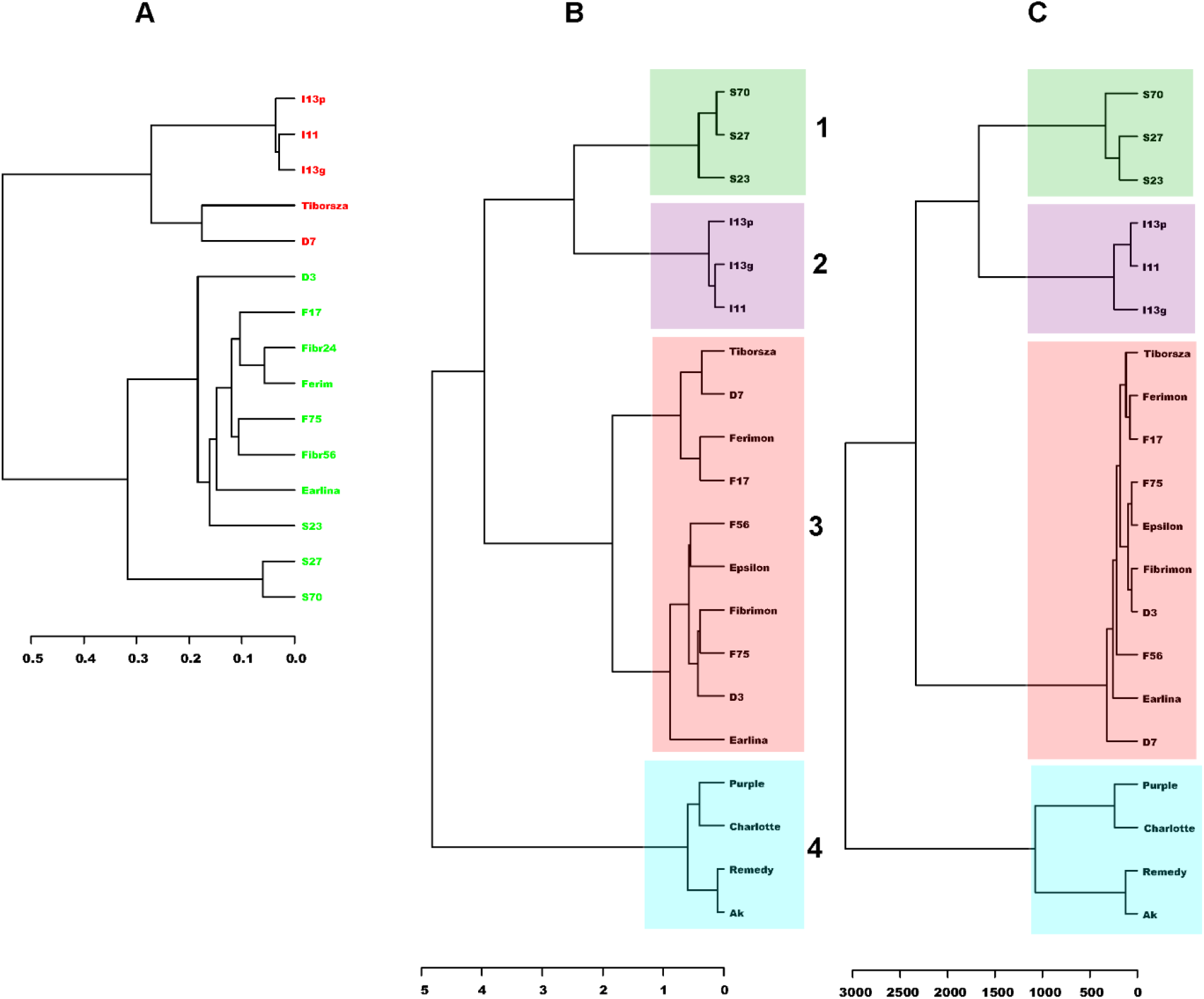
Hierarchical cluster analyses (HCA) following genetic differentiation (Fst) (**A**), the global metabolome (**B**), and main phytocannabinoids quantified (**C**). Varieties of *Cannabis*: **S27** = *Santhica 27*; **S23**= *Santhica 23*; **S70**= *Santhica 70*; **Tiborsza**= *Tiborszallasi*; **D3**= *Dioïca 3*; **D7**= *Dioïca 7*; **Ferimon**= *Ferimon*; **F17**= *Fedora 17*; **F56**= *Fibrimon F56*; **F75**= *Futura 75*; **Epsilon**= *Epsilon 68*; **Fibrimon**= *Fibrimon 24*; **Earlina**= *Earlina 8FC*; **I11**= *I11*; **I13g**= *I13g*; **I13p**= *I13p*, **AK**= *AK Silver*; **Charlotte**= *Charlotte*; **Purple**= *Purple*; **Remedy**= *Remedy*.

PCA analysis is a preliminary step in a multivariate analysis to provide an unsupervised overview of LC-MS fingerprints. An unsupervised PCA analysis on MetaboAnalyst 5.0 [44] was carried out to determine how chemotype metabolomes differ from each other, and which metabolites contribute the most to this difference (**Figure 3-A**). In our analysis, 54% of the total variance was projected in two principal component axes. As expected, replicated injections of the quality control sample (QC, made by mixing an aliquot of each extract) were grouped near the center of the plot (data not shown). Three chemotype groups were observed. On the PC1, all varieties of chemotypes III and II (hemp type) are separated from chemotype I (drug type). The differentiation between chemotypes III and II can be observed along PC2. According to the loading plot (data not shown), the separation in Factor 1 and 2 is not attributed to specific metabolites, but is a combined effect of a variety of metabolites. To validate the HCA model, we performed a supervised PLSDA analysis (Partial Least Squares Discriminant Analysis) based on the VIP (variable importance in projection) values. Overall, 47.5% of the total variance was displayed on the first two principal component axes of the PLSDA score plot (**Figure 3-B**). Samples can be separated into four subgroups/clusters: samples of the **cluster 4** were found distant of other chemotypes on the component 1, and the samples of the **cluster 1**, **2**, and **3** as three independent clusters on component 2.

**Figure 3.**
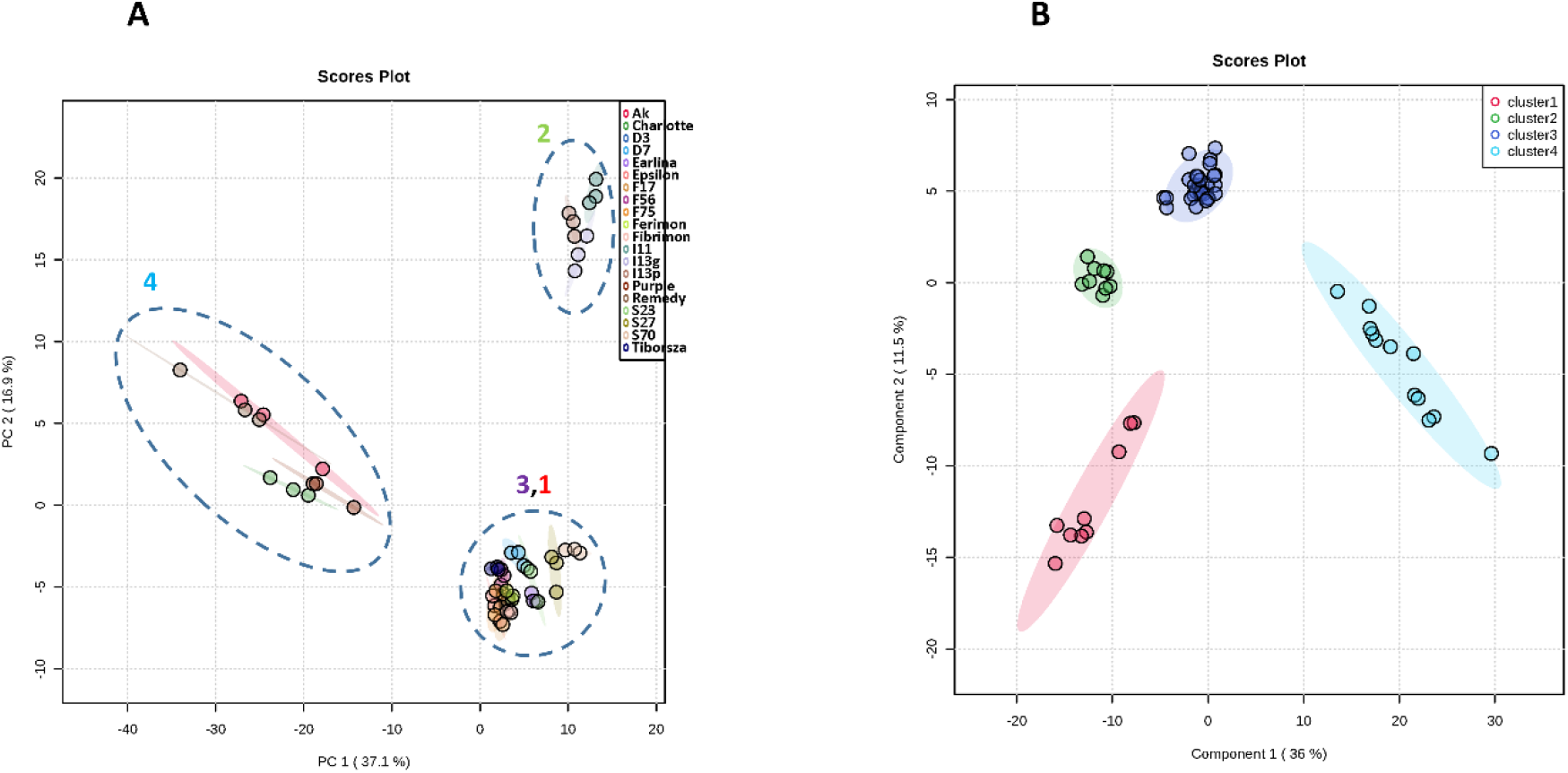
Global PCA (**A**), PLSDA analyses for clusters (**B**) for *Cannabis* chemotypes. In **A**, numbers relate to the clusters obtained in Fig 2-B.

Following, we performed a hierarchical clustering analysis using T-test/Anova to find other characteristic markers differentiating these clusters. A heat map was generated from the top 50 metabolites present in varieties separated into clusters (**Figure 4**). Based on the top 50 discriminant metabolites, this analysis results in the same four major clusters, as mentioned above. In these clusters, 24 metabolites out of 50 (48%) are phytocannabinoids; however, this type of compounds are unexpectedly not discriminant for **cluster 4.** Samples of **cluster 4** (recreative varieties) have been genetically selected to present a characteristic aroma due mostly to mono- and sesquiterpenes (*e.g.* germacrene D-type, as observed in MN, **supp info 4**). In **cluster 1**, cannabigerol (CBG) and cannabigerolic acid (CBGA) derivatives are the characteristic markers, which is consistent with the fact that this cluster groups the *Santhica* varieties, characterized by a very low concentration of THC and CBD, and significant concentrations of CBG and CBGA [32,55]. However, in **cluster 1**, other distinctive compounds are put forward, *e.g.* CBE-type (produced by photo-oxidation of CBD and CBDA) and CBCA-type compounds. Contrary to prior knowledge stating that the fiber-type hemps are characterized by CBDA [3], **cluster 3** which brings together ten fiber-type varieties show that two other cannabinoids were identified as discriminant biomarkers (Δ^4-8^-THC and a CBLA derivative). In **cluster 2**, compounds with a 3-carbon resorcinyl side-chain (called propyl phytocannabinoids or *C-3* phytocannabinoids) namely CBDVA, CBCVA, Δ^9^-THCV, and Δ^1^-THCVA are presented as the main markers. However, in this cluster, a wide diversity of phytocannabinoids is present, as we can also observe *C-5* phytocannabinoids (namely CBL, CBC, CBR, CBG, CBN, and CBM types).

**Figure 4.**
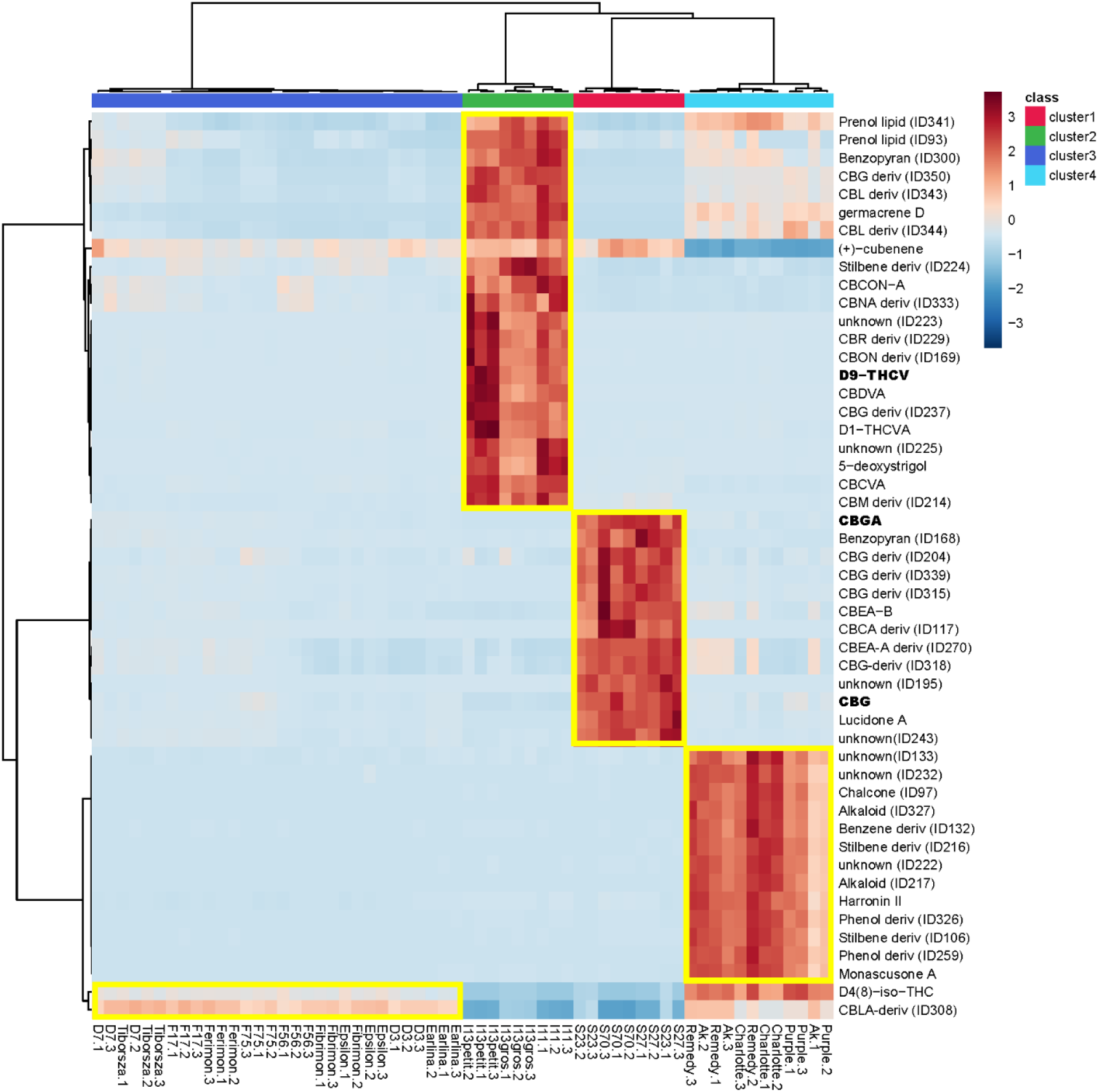
Hierarchical clustering with the heat map generated from the top 50 metabolites present in all varieties of *Cannabis*. Clusters were grouped based on the HCA analysis showed in **Figure 2**. Class of compounds were annotated using MSCleanR workflow. Phytocannabinoid standards are displayed in bold.

### 3.2. Unraveling characteristic markers of *Cannabis* varieties

To obtain specific markers for each variety within the clusters, we performed independent heatmap analyses and Anova tests. The top 25 most significant compounds are listed in **Table 1** (heatmaps in **supp info 6-9**). Among the three varieties of **cluster 1** (*Santhica* varieties with CBG as major cannabinoid), S23 appears to be very different from the two other varieties (S27 and S70) (**supp info 6**). Of the over 25 compounds discriminant of S23, 14 are phytocannabinoids including Δ^9^-THCA, Δ^8^-THC, CBN and their artifacts (CBN-1’S-hydroxy, CBNA and CBND). The CBN-type compounds result from the spontaneous oxidative aromatization of the THC-type derivatives [4]. S27 and S70 are mostly similar and share 3 markers: cannabispirenol B and two stilbenoid carexane derivatives. Discriminant markers of **cluster 2** (hemp varieties for seed production, characterized by C3 varin-type phytocannabinoids) suggest that these varieties are from different origins (**supp info 7**). Interestingly, a strigolactone derivative is identified as characteristic of two varieties. This can be related to their phenotypic aspect, which is short and much ramified. Strigolactones are phytohormones inhibiting shoot branching [56], and a change in their profile definitely has an impact on ramification. The spiran cannabispirol derivatives, like cannabispirenone A previously isolated from a Thailand drug-type *Cannabis* variety [57,58] are characteristic of I11. I13g variety showed a dozen of discriminant compounds, including CBD, which h is a decarboxylated artifact of CBDA, and a cannabisol isomer, a dimer of Δ^9^-THC with a methylene bride produced naturally in *Cannabis* plants [59]. In **cluster 3** (hemp-type varieties with CBD as major cannabinoid), 6 groups of varieties are distinguishable (**sup info 8**). As shown above by the genetic clustering (**Fig 2-A**), D7 and Tiborszallasi varieties are very close, sharing seven discriminative compounds, among them CBDVA, cannabispirenone A and a CBL derivative. A set of 6 compounds (3 of them being annotated as flavonoids, one as a fatty amide) is highly characteristic of Earlina. Three compounds are characteristic of Ferimon and F17 (including cannabistilbene I). Hildgardtol B and a phenanthrene derivative are characteristic of Epsilon. F75 is barely distinguished from other varieties but may be considered as having some similarities with Epsilon. Four compounds (one of them being annotated as the pulchelstyrene B, a cinnamylphenol) are characteristic of F56. Two stilbenoid derivatives and isocannabispiradienone appear as being discriminant for D3 and Fibrimon. These results can hardly be interpreted because chemical markers of these hemp varieties result from a massal selection applied to empirically selected populations. Phytocannabinoids can be discriminant for some varieties but not for all of them, excluding the acidic or neutral THC and CBD derivatives, which have been used as selection markers for a low content of these phytocannabinoids. In **cluster 4**, AK Silver and Remedy share the same distinctive compounds (among which Δ^9^-THCA and CBGA) (**supp info 9**). Even if Charlotte and Purple variety share Δ^9^-THC and CBD as characteristic markers, these compounds are more concentrated in Charlotte, principally Δ^9^-THC (see **Table 2**-quantification).

**Table 1.**
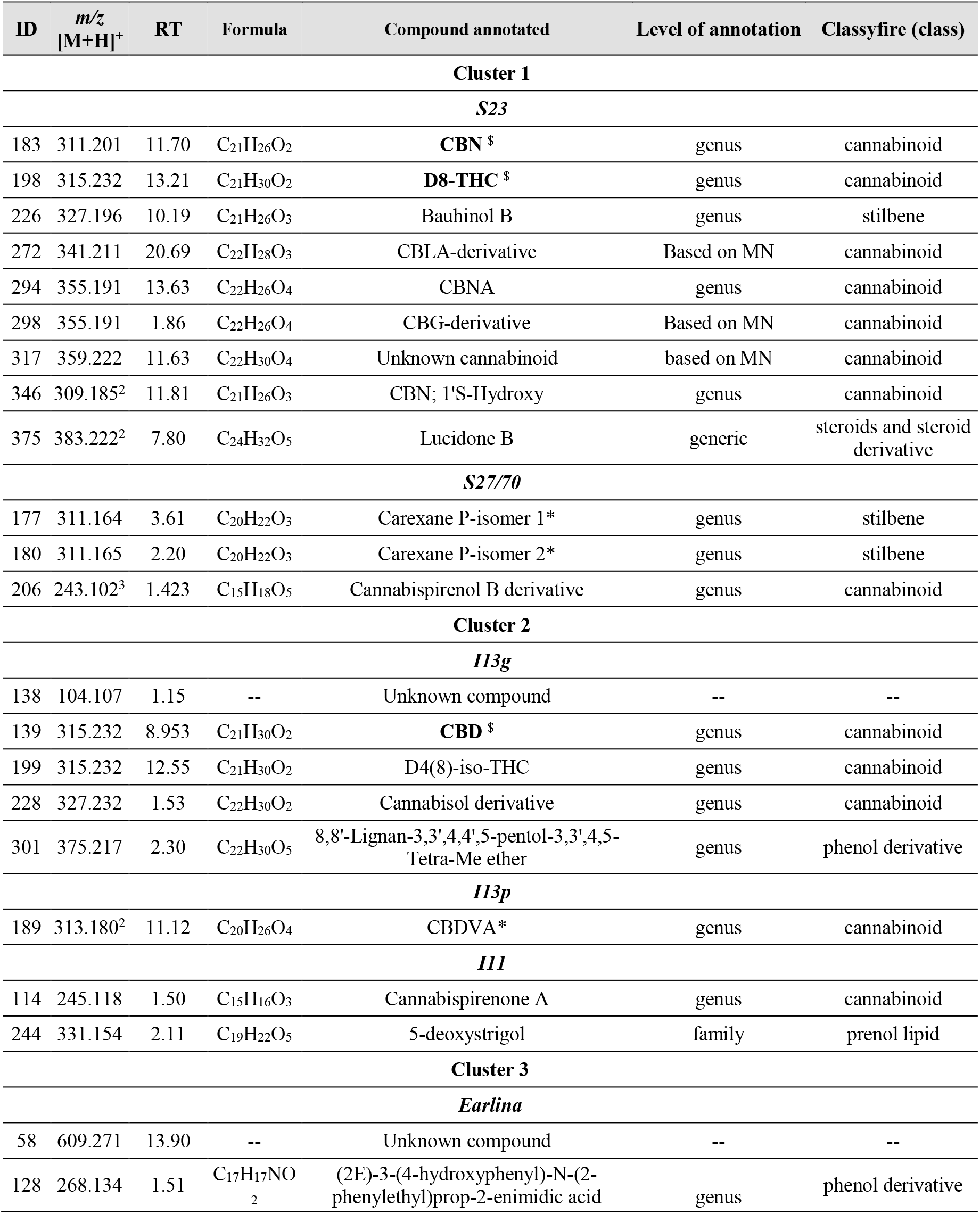
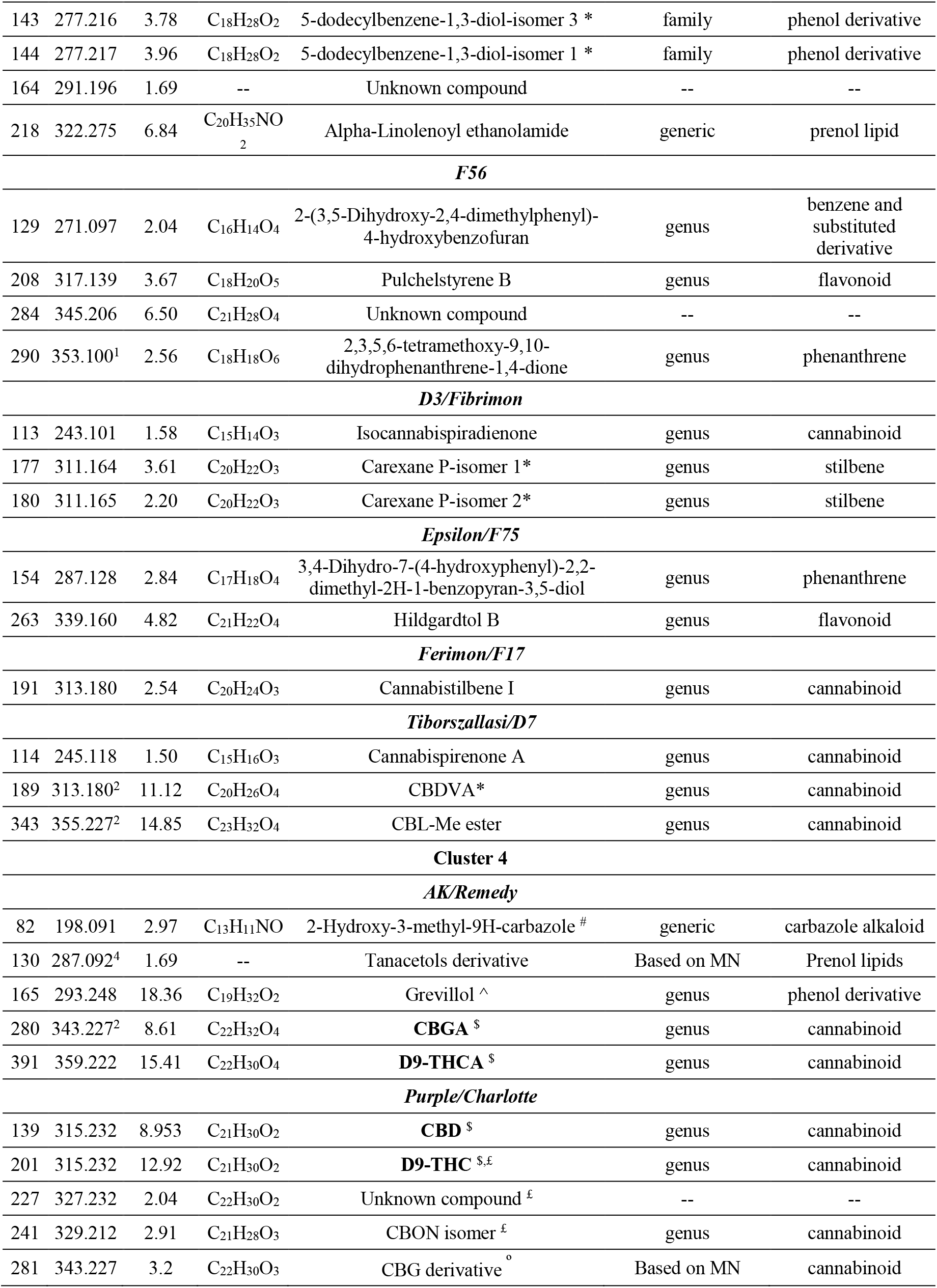
Main markers by cluster from *Cannabis* varieties based on heat map analyses. * = same annotation; ^1^ = [M+Na]^+^; ^2^ = [M+H-H_2_O]^+^; ^3^ = [M+H-2H_2_O]^+^; ^4^ = [M+K]^+^; ^$^ = standard; ^ = only in AK variety; ^#^ = only in Remedy variety; ⍛ = only in purple variety; ^£^ = only in charlotte variety. In bold: phytocannabinoids standards. MN = molecular networking.

**Table 2.** UHPLC-MS quantification of 11 phytocannabinoids from 20 *Cannabis* varieties. Results were reported as ng/mL in injected solution (% in powder), mean ± SD; n = 3.

| | $\Delta^9$-THC | $\Delta^9$-THCA | CBD | CBDA | $\Delta^8$-THC | CBG | CBGA | CBC | CBCA | $\Delta^9$-THCV | CBN |
| --- | --- | --- | --- | --- | --- | --- | --- | --- | --- | --- | --- |
| **Linearity parameters** |  |  |  |  |  |  |  |  |  |  |  |
| **Calibration curve equation** | 13950.0x + 54497.1 | 2001.8x - 16666.7 | 12003.8x - 124854.2 | 2715.1x - 191794.6 | 14995.6x + 40061.6 | 3614.5x - 186148.4 | 720.7x - 13861.0 | 4564.0x + 366666.7 | 3181.1x + 216666.7 | 11974.7x + 409756.6 | 10032.2x - 409297.1 |
| **Correlation coefficient value (R<sup>2</sup>)** | 1.0000 | 1.0000 | 0.9999 | 0.9995 | 1.0000 | 0.9997 | 0.9998 | 1.0000 | 0.9997 | 0.9997 | 0.9998 |
| **LOD (ng/mL)** | 6.7 | 24.5 | 46.9 | 219.5 | 5.0 | 164.9 | 120.8 | 187.3 | 275.0 | 109.6 | 128.4 |
| **LOQ (ng/mL)** | 19.7 | 83.7 | 165.1 | 759.0 | 14.8 | 569.7 | 410.2 | 604.4 | 900.0 | 342.6 | 443.8 |
| **Sample quantification (ng/mL in injected solution (% in powder), mean ± SD)** |  |  |  |  |  |  |  |  |  |  |  |
| **AK Silver** | 116 ± 19<br>(0.006 ± 0.001) | 32066 ± 5274<br>(1.603 ± 0.264) | 85409 ± 9776<br>(4.270 ± 0.489) | 290696 ± 25319<br>(14.535 ± 1.266) | 10387 ± 1854<br>(0.519 ± 0.093) | 3380 ± 717<br>(0.169 ± 0.036) | 22324 ± 4427<br>(1.116 ± 0.221) | 6884 ± 654<br>(0.344 ± 0.033) | 39288 ± 5538<br>(1.964 ± 0.277) | <LOD | 991 ± 230<br>(0.050 ± 0.012) |
| **Charlotte** | 941 ± 118<br>(0.047 ± 0.006) | 2116 ± 495<br>(0.106 ± 0.025) | 144968 ± 7134<br>(7.248 ± 0.357) | 147590 ± 19191<br>(7.380 ± 0.960) | 9393 ± 992<br>(0.470 ± 0.050) | 1673 ± 135<br>(0.050 ± 0.007) | 1693 ± 181<br>(0.085 ± 0.009) | 11887 ± 1170<br>(0.594 ± 0.058) | 12313 ± 2489<br>(0.616 ± 0.124) | <LOQ | 1989 ± 270<br>(0.099 ± 0.013) |
| **Purple** | 715 ± 73<br>(0.036 ± 0.004) | 7344 ± 1040<br>(0.367 ± 0.052) | 140928 ± 1717<br>(7.046 ± 0.086) | 187738 ± 2401<br>(9.387 ± 0.120) | 17086 ± 1837<br>(0.854 ± 0.092) | 5193 ± 1033<br>(0.260 ± 0.052) | 5786 ± 620<br>(0.289 ± 0.031) | 15620 ± 1732<br>(0.781 ± 0.087) | 25653 ± 3842<br>(1.283 ± 0.192) | <LOD | 1343 ± 149<br>(0.067 ± 0.007) |
| **Remedy** | 141 ± 51<br>(0.007 ± 0.003) | 40302 ± 11370<br>(2.015 ± 0.569) | 96951 ± 29408<br>(4.848 ± 1.470) | 302006 ± 37892<br>(15.100 ± 1.895) | 13180 ± 5571<br>(0.659 ± 0.279) | 5118 ± 2103<br>(0.256 ± 0.105) | 26266 ± 10919<br>(1.313 ± 0.546) | 8875 ± 3534<br>(0.444 ± 0.177) | 44744 ± 15337<br>(2.237 ± 0.767) | <LOD | 1066 ± 448<br>(0.053 ± 0.022) |
| **D3** | 47 ± 6<br>(0.002 ± 0.000) | 3299 ± 271<br>(0.165 ± 0.014) | 37510 ± 4630<br>(1.875 ± 0.231) | 102088 ± 13455<br>(5.104 ± 0.673) | 2594 ± 43<br>(0.130 ± 0) | 1033 ± 31<br>(0.052 ± 0.002) | 2050 ± 232<br>(0.103 ± 0.012) | 2887 ± 196<br>(0.144 ± 0.010) | 8299 ± 506<br>(0.415 ± 0.025) | <LOD | 557 ± 63<br>(0.028 ± 0.003) |
| **D7** | 34 ± 17<br>(0.002 ± 0.001) | 18150 ± 3829<br>(0.908 ± 0.191) | 23727 ± 10174<br>(1.186 ± 0.509) | 158525 ± 39631<br>(7.926 ± 1.982) | 5078 ± 2341<br>(0.254 ± 0.117) | 1759 ± 835<br>(0.088 ± 0.042) | 13110 ± 4507<br>(0.656 ± 0.225) | 2125 ± 851<br>(0.106 ± 0.043) | 16246 ± 6595<br>(0.812 ± 0.330) | <LOD | <LOQ |
| **Earlina** | 54 ± 1<br>(0.003 ± 0.000) | 1775 ± 160<br>(0.089 ± 0.008) | 29158 ± 1224<br>(1.458 ± 0.061) | 69743 ± 1338<br>(3.487 ± 0.067) | 1783 ± 80<br>(0.089 ± 0.004) | 1271 ± 24<br>(0.064 ± 0.001) | 2922 ± 353<br>(0.146 ± 0.018) | 3028 ± 181<br>(0.151 ± 0.009) | 5965 ± 169<br>(0.298 ± 0.008) | <LOD | <LOQ |
| **Epsilon** | 62 ± 25<br>(0.003 ± 0.001) | 4792 ± 2036<br>(0.240 ± 0.102) | 41796 ± 13520<br>(2.090 ± 0.676) | 125640 ± 26148<br>(6.282 ± 1.307) | 3481 ± 1469<br>(0.174 ± 0.073) | 1424 ± 490<br>(0.071 ± 0.024) | 4612 ± 2008<br>(0.231 ± 0.100) | 3446 ± 1456<br>(0.172 ± 0.073) | 11243 ± 5441<br>(0.562 ± 0.272) | <LOD | 537 ± 196<br>(0.027 ± 0.010) |
| **Ferimon** | 36 ± 5<br>(0.002 ± 0.000) | 5939 ± 599<br>(0.297 ± 0.030) | 28286 ± 769<br>(1.414 ± 0.038) | 144063 ± 4446<br>(7.203 ± 0.222) | 2730 ± 200<br>(0.136 ± 0.010) | 1790 ± 90<br>(0.089 ± 0.005) | 9119 ± 1615<br>(0.456 ± 0.081) | 2605 ± 69<br>(0.130 ± 0.003) | 13563 ± 1837<br>(0.678 ± 0.092) | <LOD | <LOQ |
| **Fibrimon** | 62 ± 4<br>(0.003 ± 0.000) | 4242 ± 393<br>(0.212 ± 0.020) | 32133 ± 3425<br>(1.607 ± 0.171) | 110474 ± 9044<br>(5.524 ± 0.452) | 2565 ± 281<br>(0.128 ± 0.014) | 1008 ± 16<br>(0.050 ± 0.001) | 3023 ± 43<br>(0.151 ± 0.002) | 3903 ± 215<br>(0.195 ± 0.011) | 11904 ± 844<br>(0.595 ± 0.042) | <LOD | 487 ± 17<br>(0.024 ± 0.001) |
| **F17** | 46 ± 20<br>(0.002 ± 0.001) | 4481 ± 2638<br>(0.224 ± 0.132) | 29647 ± 11918<br>(1.482 ± 0.596) | 122112 ± 39417<br>(6.106 ± 1.971) | 2840 ± 1324<br>(0.142 ± 0.066) | 1833 ± 793<br>(0.092 ± 0.040) | 8822 ± 5449<br>(0.441 ± 0.272) | 3306 ± 1489<br>(0.165 ± 0.074) | 13188 ± 7568<br>(0.659 ± 0.378) | <LOD | <LOQ |
| **F56** | 54 ± 29<br>(0.003 ± 0.001) | 15860 ± 515<br>(0.793 ± 0.026) | 37256 ± 13884<br>(1.863 ± 0.694) | 106675 ± 23241<br>(5.334 ± 1.162) | 8117 ± 3986<br>(0.406 ± 0.199) | 1388 ± 630<br>(0.069 ± 0.031) | 5602 ± 2231<br>(0.280 ± 0.112) | 3090 ± 1624<br>(0.155 ± 0.081) | 9703 ± 3136<br>(0.485 ± 0.157) | <LOD | 1566 ± 764<br>(0.078 ± 0.038) |
| **F75** | 71 ± 26<br>(0.004 ± 0.001) | 3949 ± 1422<br>(0.197 ± 0.071) | 43767 ± 18352<br>(2.188 ± 0.918) | 112857 ± 30611<br>(5.643 ± 1.531) | 2965 ± 1062<br>(0.148 ± 0.053) | 2941 ± 1427<br>(0.147 ± 0.071) | 8964 ± 4582<br>(0.448 ± 0.229) | 4036 ± 1455<br>(0.202 ± 0.073) | 10033 ± 4855<br>(0.502 ± 0.243) | <LOD | 601 ± 254<br>(0.030 ± 0.013) |
| **I11** | <LOQ | 53728 ± 16170<br>(2.686 ± 0.808) | 801 ± 368<br>(0.040 ± 0.018) | 8220 ± 3343<br>(0.411 ± 0.167) | 8902 ± 4045<br>(0.445 ± 0.202) | <LOQ | 1117 ± 569<br>(0.056 ± 0.028) | 698 ± 419<br>(0.035 ± 0.021) | 8376 ± 3880<br>(0.419 ± 0.194) | 4848 ± 0.242<br>(2307 ± 0.115) | <LOQ |
| **I13gros** | <LOQ | 27328 ± 3244<br>(1.366 ± 0.162) | 1827 ± 114<br>(0.091 ± 0.006) | 22512 ± 777<br>(1.126 ± 0.039) | 6002 ± 367<br>(0.300 ± 0.018) | <LOQ | 792 ± 148<br>(0.040 ± 0.007) | 682 ± 63<br>(0.034 ± 0.003) | 7288 ± 1210<br>(0.364 ± 0.061) | 3302 ± 111<br>(0.165 ± 0.006) | <LOQ |
| **I13petit** | <LOQ | 48151 ± 2710<br>(2.408 ± 0.135) | 661 ± 59<br>(0.033 ± 0.003) | 9161 ± 1659<br>(0.458 ± 0.083) | 7306 ± 548<br>(0.365 ± 0.027) | <LOQ | 1132 ± 74<br>(0.057 ± 0.004) | <LOQ | 8223 ± 737<br>(0.411 ± 0.037) | 6779 ± 591<br>(0.339 ± 0.030) | <LOQ |
| **S23** | 44 ± 23<br>(0.002 ± 0.001) | 1404 ± 514<br>(0.070 ± 0.026) | 8415 ± 854<br>(0.421 ± 0.043) | 44425 ± 342<br>(2.221 ± 0.017) | 1084 ± 484<br>(0.054 ± 0.024) | 16492 ± 7565<br>(0.825 ± 0.378) | 131988 ± 42368<br>(6.599 ± 2.118) | 2529 ± 1109<br>(0.126 ± 0.055) | 5737 ± 2409<br>(0.287 ± 0.120) | <LOD | <LOD |
| **S27** | 66 ± 32<br>(0.003 ± 0.002) | 328 ± 143<br>(0.016 ± 0.007) | 1698 ± 812<br>(0.085 ± 0.041) | 11361 ± 5199<br>(0.568 ± 0.260) | 170 ± 89<br>(0.008 ± 0.004) | 12024 ± 6055<br>(0.601 ± 0.303) | 123691 ± 36527<br>(6.185 ± 1.826) | 4048 ± 1641<br>(0.202 ± 0.082) | 4159 ± 2023<br>(0.208 ± 0.101) | <LOD | <LOD |
| **S70** | 40 ± 5<br>(0.002 ± 0.000) | <LOQ | 772 ± 34<br>(0.039 ± 0.002) | 3094 ± 351<br>(0.155 ± 0.018) | 70 ± 6<br>(0.003 ± 0.000) | 7642 ± 1511<br>(0.382 ± 0.076) | 78910 ± 8265<br>(3.945 ± 0.413) | 1877 ± 134<br>(0.094 ± 0.007) | 1825 ± 427<br>(0.091 ± 0.021) | <LOD | <LOD |
| **Tiborszallasi** | 25 ± 1<br>(0.001 ± 0.000) | 12556 ± 2338<br>(0.628 ± 0.117) | 22429 ± 765<br>(1.121 ± 0.038) | 116672 ± 6549<br>(5.834 ± 0.327) | 5704 ± 399<br>(0.285 ± 0.020) | 1544 ± 111<br>(0.077 ± 0.006) | 9795 ± 523<br>(0.490 ± 0.026) | 1877 ± 134<br>(0.094 ± 0.007) | 10394 ± 814<br>(0.520 ± 0.041) | <LOD | <LOQ |

### 3.3. Classification based on targeted phytocannabinoids quantification

Dozens of methods are used to quantify phytocannabinoids, based on many techniques: LC-HRMS [60], LC-DAD [61], HPTLC [62], GC-FID [30,63], GC-MS [64], Nuclear Magnetic Resonance (NMR) [65], and Triple quadrupole Multiple Reaction Monitoring (QqQ-MRM) [66], among others. Techniques and methods used in *Cannabis* quality control monographs differ between countries. Phytocannabinoids as a class of compounds are characterized by the occurrence of many isomers and the labile nature of the native acidic forms. Such complexity is well depicted in the molecular network of *Cannabis* extracts (**Figure 1**). As a consequence, the quantification of phytocannabinoids is usually based only on the quantification of their major compounds. We quantified 11 phytocannabinoids using LCMS (**Table 2** and **sup info 10-20**). Quantitative results match with markers identified in **Figure 4**, notably for Δ^9^-THCV (detected only in **cluster 2**) and CBG-CBGA mainly present in **cluster 1** (**sup info 14**, **15**). It is important to highlight that acidic phytocannabinoids are non-enzymatically decarboxylated into their corresponding neutral forms, which occur both within the plant and, to a much larger extent, upon heating after harvesting [67]. Therefore, acidic and neutral forms are related to a certain extent, depending on the temperature of drying. All samples studied were dried at 40°C, except for samples from **cluster 4** which were dried at 80°C. Δ^9^-THCA, the native form of Δ^9^-THC, is elevated in **cluster 2** (chemotype II, intermediate type, seed-producing varieties) and surprisingly still present in **cluster 3** (chemotype III, fiber-type) (**supp info 19**). Unsupervised PLSDA based on the results of the quantification of 11 phytocannabinoids showed that **clusters 2**, **3,** and **4** are grouped on component 2, and **cluster 1** on component 1 (**Figure 5-A**). 62.9% of the total variance was displayed on the first two principal component axes of the PLSDA score plot. To identify the most important metabolites allowing discrimination between the clusters, we performed a supervised PLSDA based on the variable importance in projection (VIP) values. Thus, a metabolite with a VIP > 1 is regarded as significantly discriminant. CBD, CBDA and CBGA are the overall main discriminant quantified phytocannabinoids (**Figure 5-B**). Some varieties presented as drug-type can be assimilated to fiber-types, *e.g. AK* and *Remedy* samples (**Figure 5 A** and **C**), although their phytocannabinoids content is higher (**Table 2**). Such drug-type varieties have been selected to produce significant amount of CBD but only moderate amounts of THC in order to be considered as “light cannabis” in some countries (close to 0,2 % Δ^9^-THC), thus these recreative varieties are barely distinguished from hemp varieties when clustering is based only on main phytocannabinoids. Hemp varieties have been specifically selected to have a phytocannabinoids biosynthetic machinery which is significantly inhibited. Samples of **cluster 2** stand out because of their specific expression of Δ^9^-THCV, and also because of a significant concentration of Δ^9^-THCA. As observed above, such varieties are used for seed production outside EU (mostly in China) but have not been subject to an efficient selection to reduce Δ^9^-THC content. As expected, **cluster 1** (hemp varieties with CBG as major cannabinoid) are characterized by CBG and CBGA.

**Figure 5.**
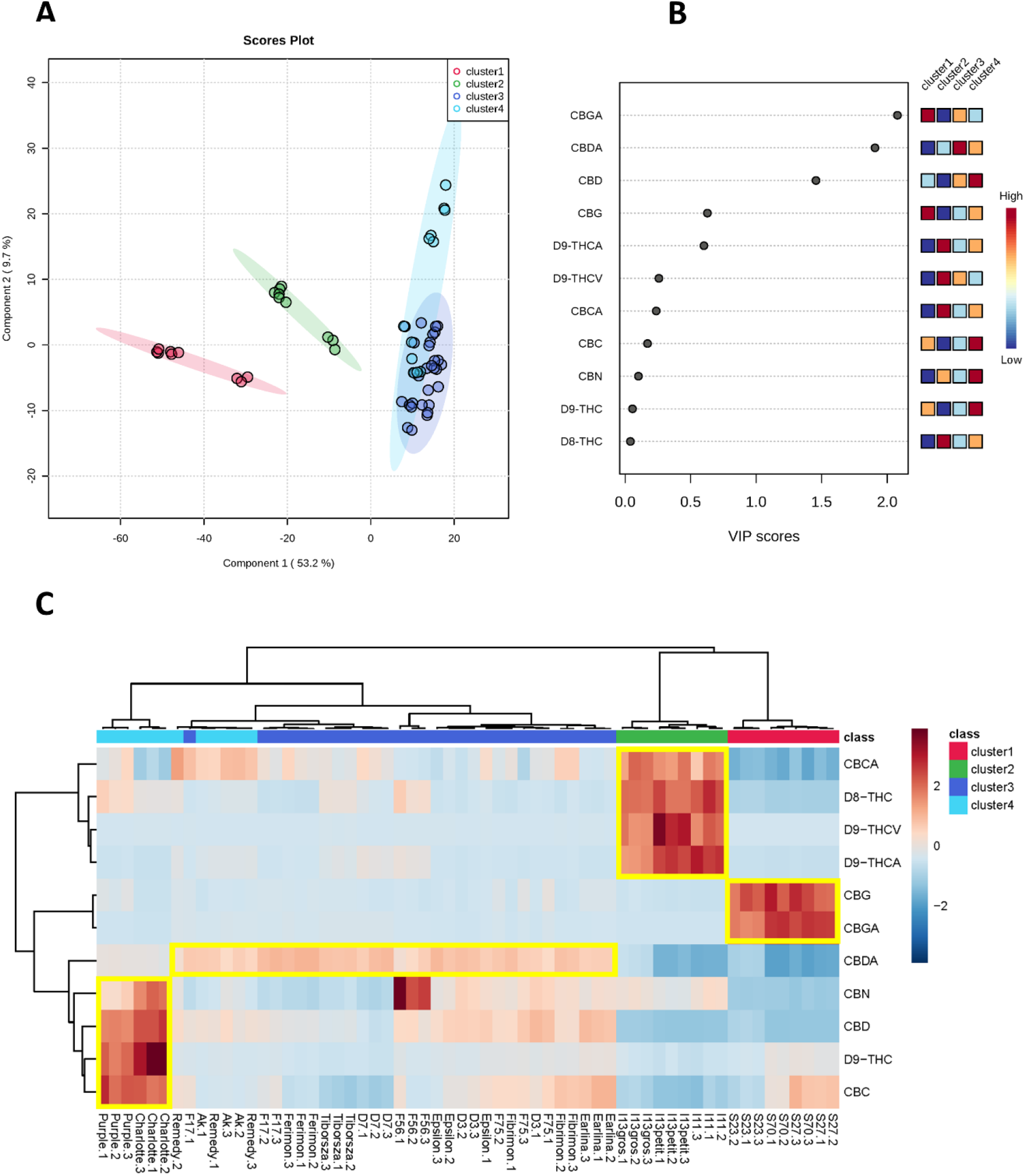
Statistical analyses based of quantitative data on 11 phytocannabinoids: PLSDA based on average concentration (ng/mL) for each phytocannabinoid by cluster (obtained in **Figure 2-C**) (**A**); PLSDA-VIP projection by cluster (**B**); hierarchical clustering analysis with the heat-map for phytocannabinoid standards by clusters (**C**).

An overall analysis of the data shows that *Cannabis* varieties are more efficiently discriminated and characterized by untargeted cannabinomics analysis (metabotyping) rather than quantitation of major phytocannabinoids. In this sense, the performance measure (Q2 and R2) were compared between metabolomic and quantification models on PLSDA analysis, suggesting that the metabotyping represents a most robust model (0.39 vs. 0.77, 0.26 vs. 0.75 on comp1 and 0.50 vs. 0.96, 0.45 vs. 0.94 on comp2 respectively).

## 4. Conclusions

Literature on *Cannabis* is expanding quickly, but is still mostly focused on their pharmacologically active compounds Δ^9^-THC and CBD. The metabolomics approach is a powerful tool to unravel the complexities of the actual entire metabolome. Cannabinomics must therefore be understood as an approach including the analyses not only of phytocannabinoids but also other compounds including terpenes, stilbenes, flavonoids, etc. In this work, cannabinomics contributed to deciphering the main discriminant metabolites in 20 *Cannabis* varieties. Among the 13 fiber-type varieties, we identified characteristic markers allowing an efficient differentiation beyond phytocannabinoids only. Varieties of the intermediate chemotype used for seed production (“I” varieties) are identified as mostly producing *C-3* phytocannabinoids. This workflow can be used as a complementary tool to be used along with genotyping with the aim to differentiate *C. sativa* varieties and to allow the identification of minor compounds playing a key role as markers of differentiation. This workflow could offer a shorter road to creating chemotypes and phenotypes that meet the demand of production needs for material, food and medicinal purposes. In the medicinal field, cannabinomics could facilitate improved breeding to create *Cannabis* varieties with greater expression of minor phytocannabinoids of medicinal value. Cannabinomics should also be used as a systematic basis for studies correlating chemical profile and therapeutical effects, as it allows to properly address the entourage effect.

## Supporting information

supp info 1 - table of varieties

supp info2_SSR markers

supp info3_Metadata

supp info4_mol network_cluster

supp info5_genotyping-structure

supp info6_heatmap-cluster1

supp info6_heatmap-cluster2

supp info6_heatmap-cluster3

supp info6_heatmap-cluster4

supp info10_CBC-violin

supp info11_CBCA-violin

supp info12_CBD-violin

supp info13_CBDA-violin

supp info14_CBG-violin

supp info15_CBGA-violin

supp info16_CBN-violin

supp info17_D8-THC-violin

supp info18_D9-THC-violin

supp info19_D9-THCA-violin

supp info20_D9-THCV-violin

## Author’s contribution

Conceptualization: PGV-O, AM and GM; methodology: PGV-O, AM, GM, SF and SB; chemical data acquisition/curation: PGVO and GM; biological data curation: MB and FM; investigation: PGV-O, GM, FM, MB, SF, SB and AM. PGVO wrote the original draft and all the authors contributed to the final manuscript.

## Acknowledgments

The authors wish to acknowledge HEMP-it ADN for the postdoctoral fellowship of PGV-O (CNRS UMR 8076 BioCIS). PGV-O was also supported by a postdoctoral fellowship (contract number 04077858) from the French National Research Institute for Sustainable Development (IRD), UMR 152 PHARMADEV, IRD-UPS, Toulouse, France. The authors are grateful to Bruno Figadère (CNRS UMR 8076 BioCIS, Châtenay-Malabry); Justine Chervin and Virginie Puech-Pagès (MetaboHUB-MetaToul-Agromix, LRSV, Auzeville-Tolosane); Guillaume Cabanac (IRIT, Toulouse); Claire Thouminot and Christophe Fevrier (HEMP-it ADN, Beaufort-en-Anjou). The authors are also grateful to Alice Gadea (UMR 152 PHARMADEV, IRD-UPS, Toulouse, France), Alice Kan, and Joshua Lee Halford (Department of Biochemistry, UCR, Riverside, USA) for their important critical discussions.

