## Supplementary material for "Cannabinoids vs. whole metabolome: relevance of cannabinomics in analyzing *Cannabis* varieties": supp info 1 - table of varieties

| Codes/ID | Name | Drying | Cultivation type | Main native cannabinoids | chemotype | Use | Geographical origin | Comment |
|---|---|---|---|---|---|---|---|---|
| **AK** | AK Silver | 80°C oven | Hydroponic | THC/CBD | I | Recreative | Austria | Legal in some EU countries |
| **Charlotte** | Charlotte | 80°C oven | Hydroponic | THCA/CBDa | I | Recreative | Italy | Legal in some EU countries |
| **D3** | Dioïca3 | 40°C oven | Open field | CBDA | III | Fibers | France | EU list* |
| **D7** | Dioïca7 | 40°C oven | Open field | CBDA | III | Fibers | France | EU list* |
| **Earlina** | Earlina 8FC | 40°C oven | Open field | CBDA | III | Fibers | France | EU list* |
| **Epsilon** | Epsilon 68 | 40°C oven | Open field | CBDA | III | Fibers | France | EU list* |
| **Ferimon** | Ferimon | 40°C oven | Open field | CBDA | III | Fibers | France | EU list* |
| **Fibrimon** | Fibrimon | 40°C oven | Open field | CBDA | III | Fibers | France | EU list* |
| **F17** | Fedora 17 | 40°C oven | Open field | CBDA | III | Fibers | France | EU list* |
| **F56** | F56 | 40°C oven | Open field | CBDA | III | Fibers | France | EU list* |
| **F75** | Futura 75 | 40°C oven | Open field | CBDA | III | Fibers | France | EU list* |
| **I11** | I11 chinois | 40°C oven | Open field | THCVA/CBDVA | II | Seeds | China | Hempit AND Germplasm database |
| **I13g** | I13 chinois gros | 40°C oven | Open field | THCVA | II | Seeds | China | Hempit AND Germplasm database |
| **I13p** | I13 chinois petit | 40°C oven | Open field | THCVA | II | Seeds | China | Hempit AND Germplasm database |
| **Purple** | Purple | 80°C oven | Hydroponic | THCVA | I | Recreative | Italy | Legal in some EU countries |
| **Remedy** | Remedy | 80°C oven | Hydroponic | THCA/CBDA | I | Recreative | Austria | Legal in some EU countries |
| **Tiborsza** | Tiborszallasi | 40°C oven | Open field | THCA/CBDA | III | Fibers | France | EU list* |
| **S23** | Santhica 23 | 40°C oven | Open field | CBGA | III | Fibers | France | EU list* |
| **S27** | Santhica 27 | 40°C oven | Open field | CBGA | III | Fibers | France | EU list* |
| **S70** | Santhica 70 | 40°C oven | Open field | CBGA | III | Fibers | France | EU list* |

*varieties are on the EU list of authorized hemp varieties.
