## Supplementary figures and images for "Cannabinoids vs. whole metabolome: relevance of cannabinomics in analyzing *Cannabis* varieties"

### supp info4_mol network_cluster

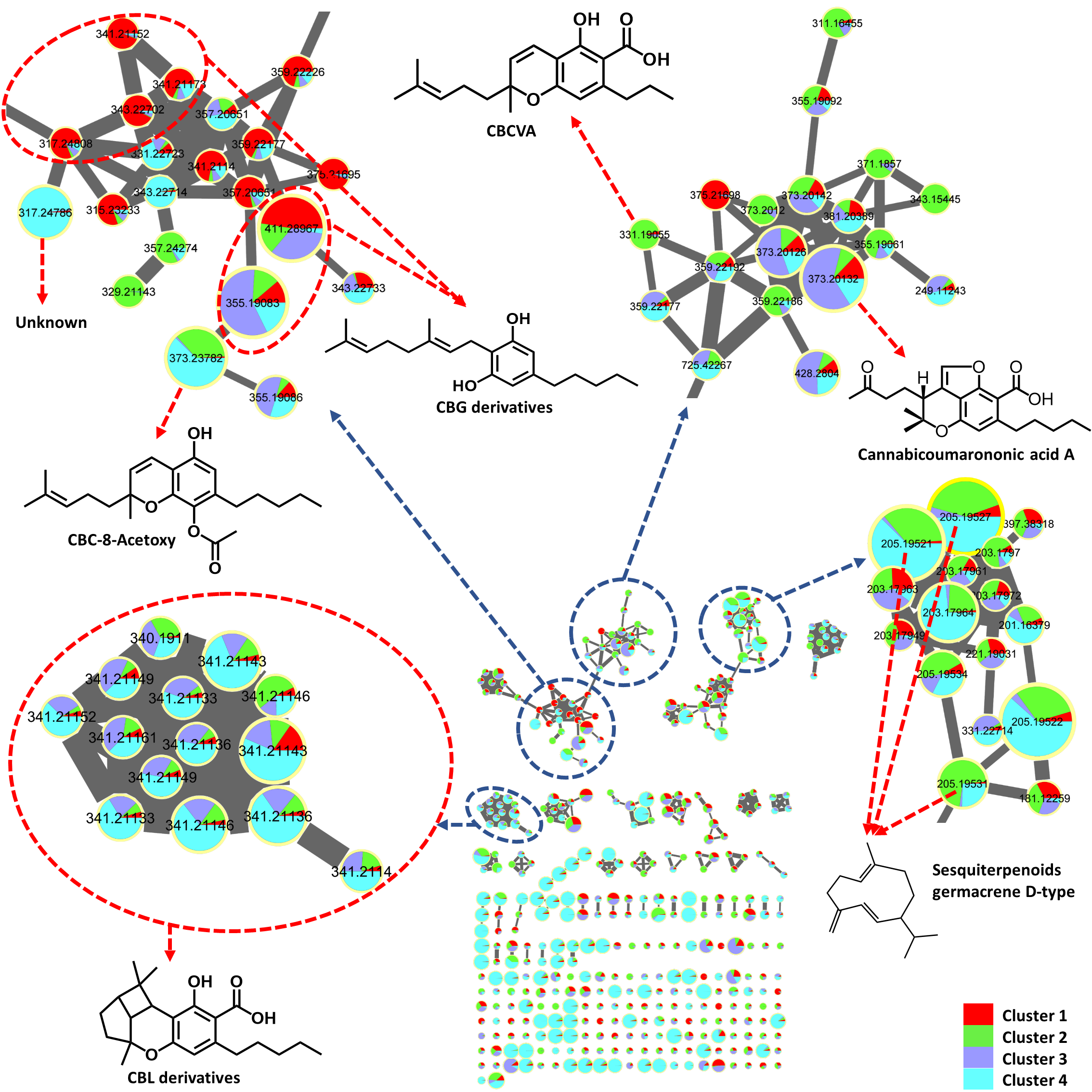

### supp info6_heatmap-cluster1

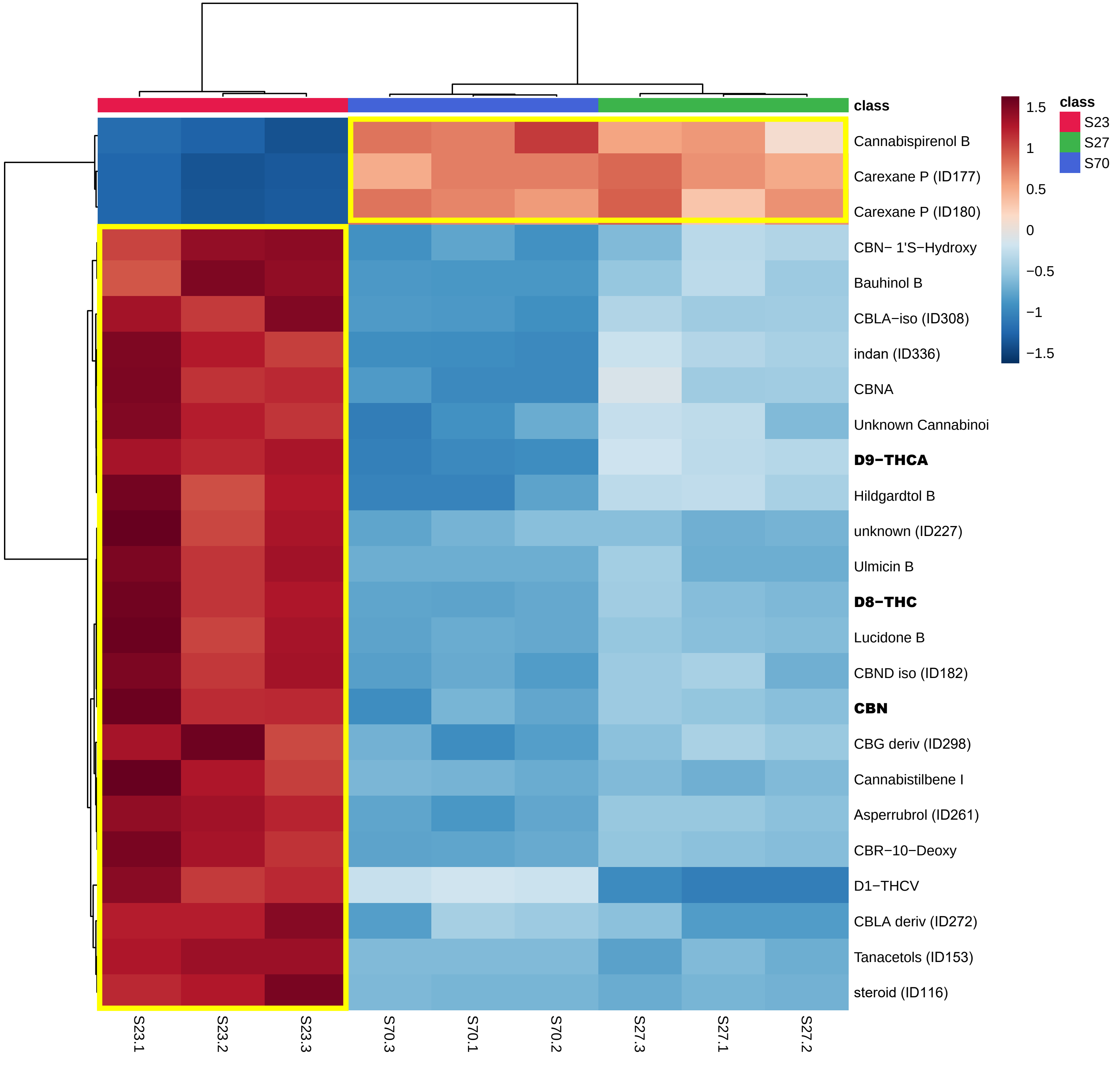

### supp info6_heatmap-cluster2

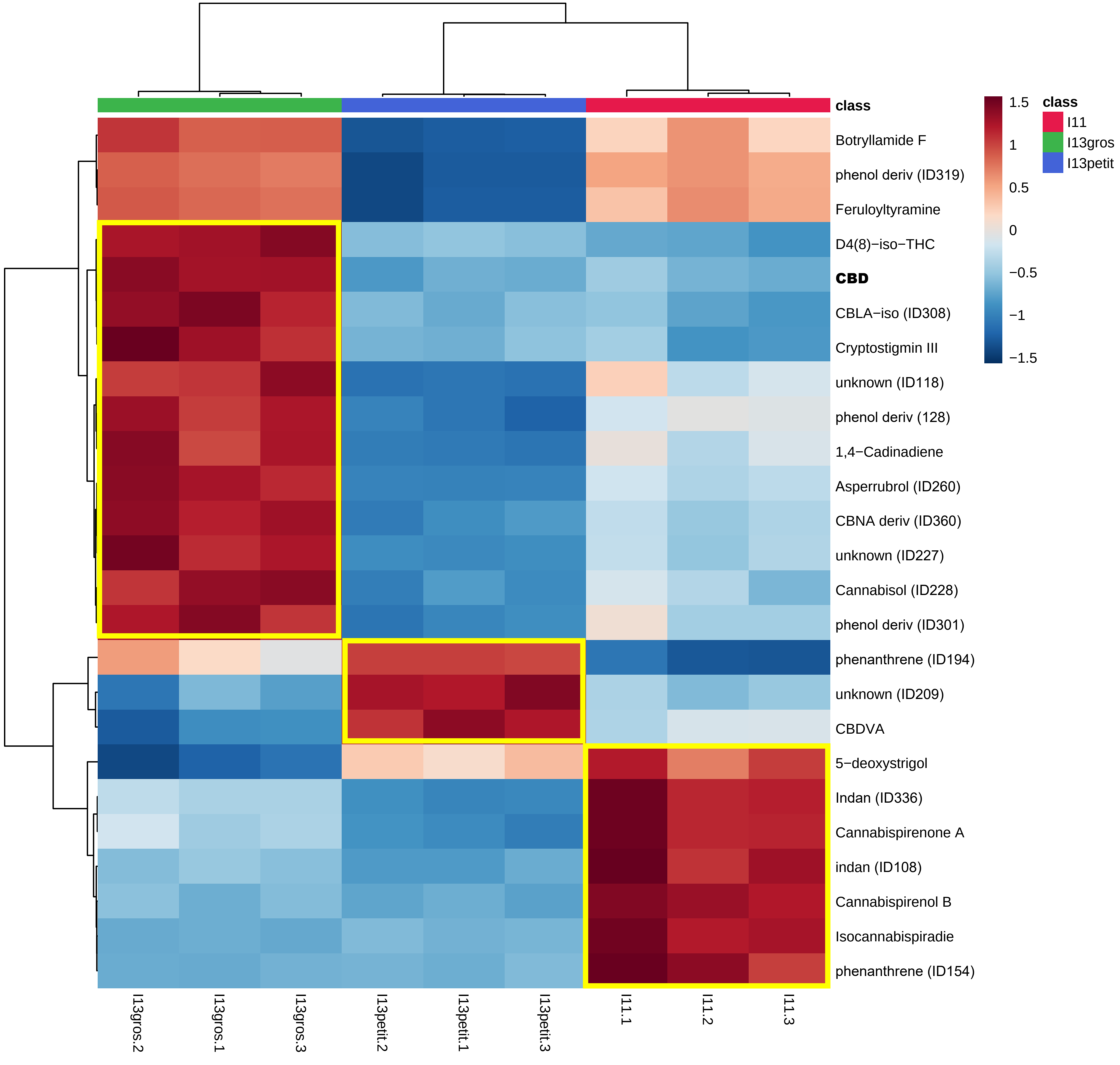

### supp info6_heatmap-cluster3

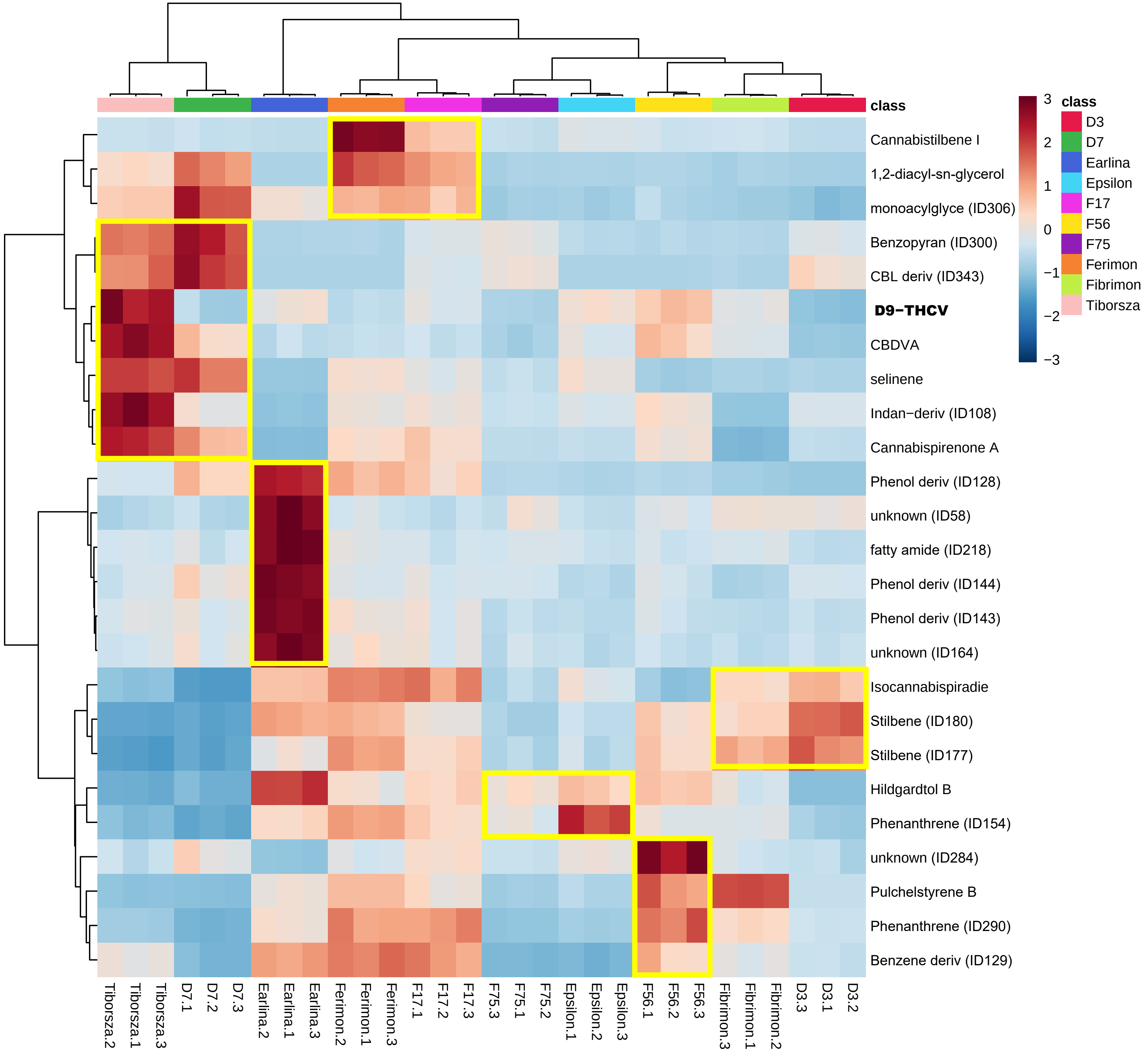

### supp info6_heatmap-cluster4

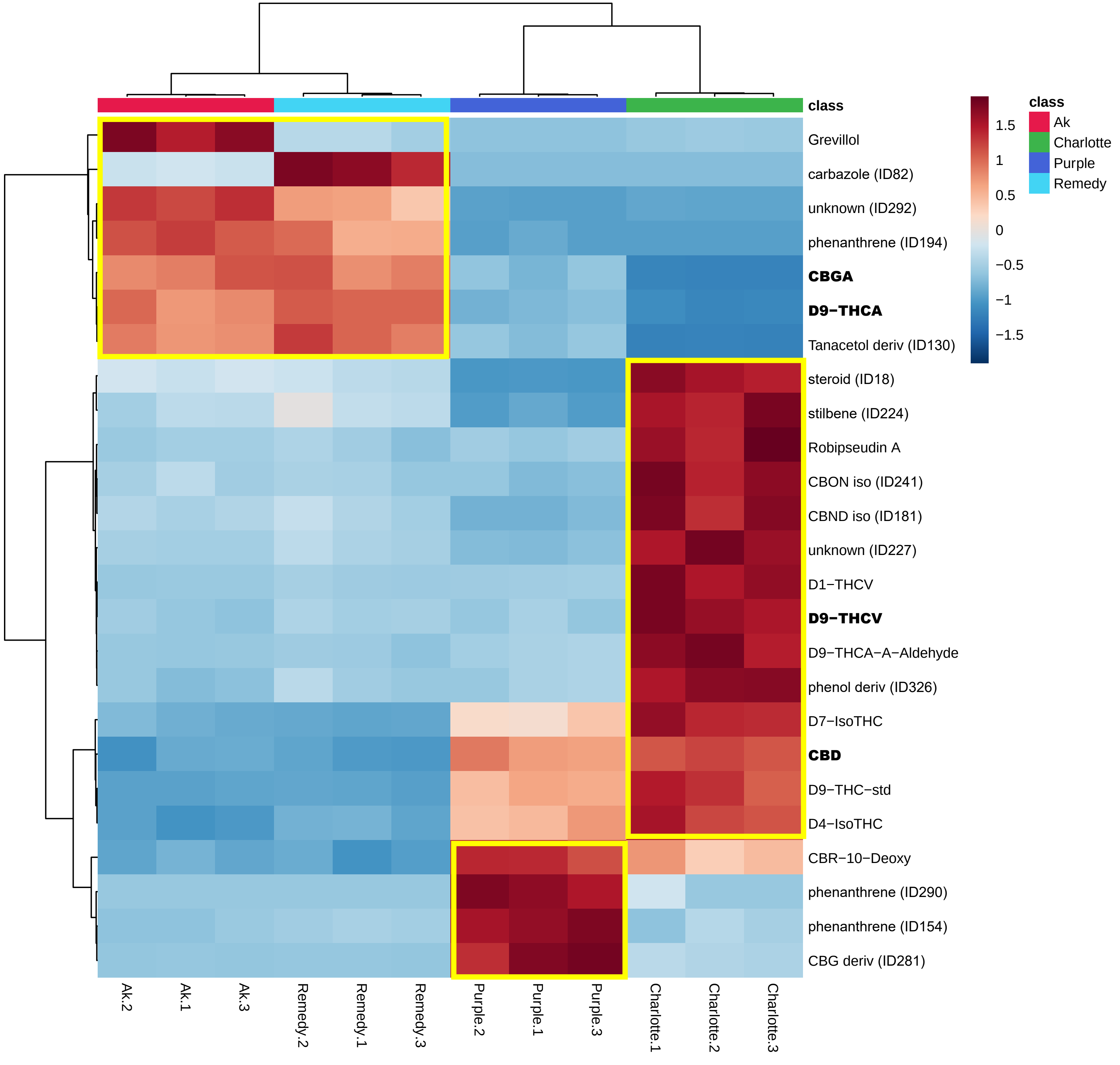

### supp info10_CBC-violin

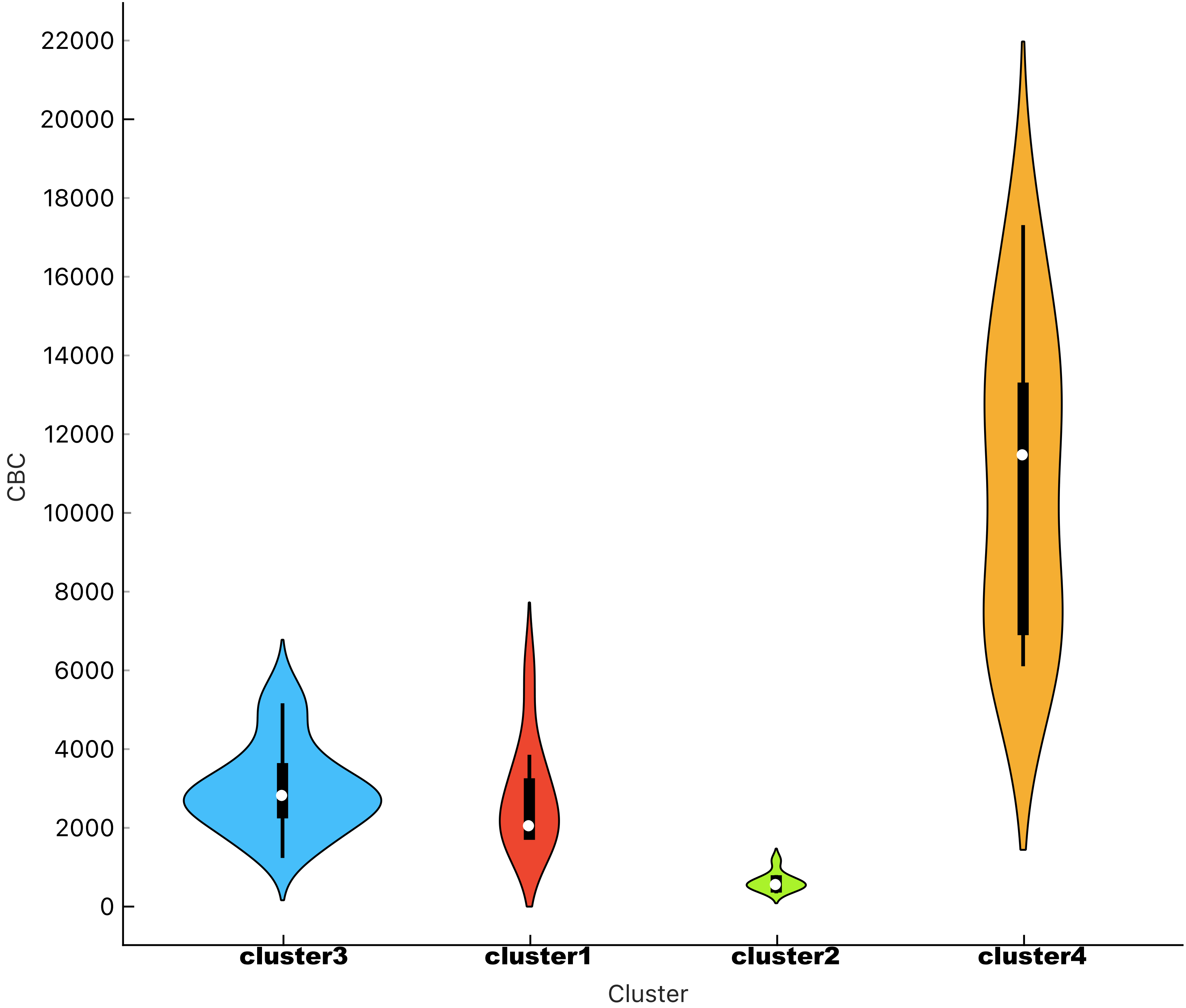

### supp info11_CBCA-violin

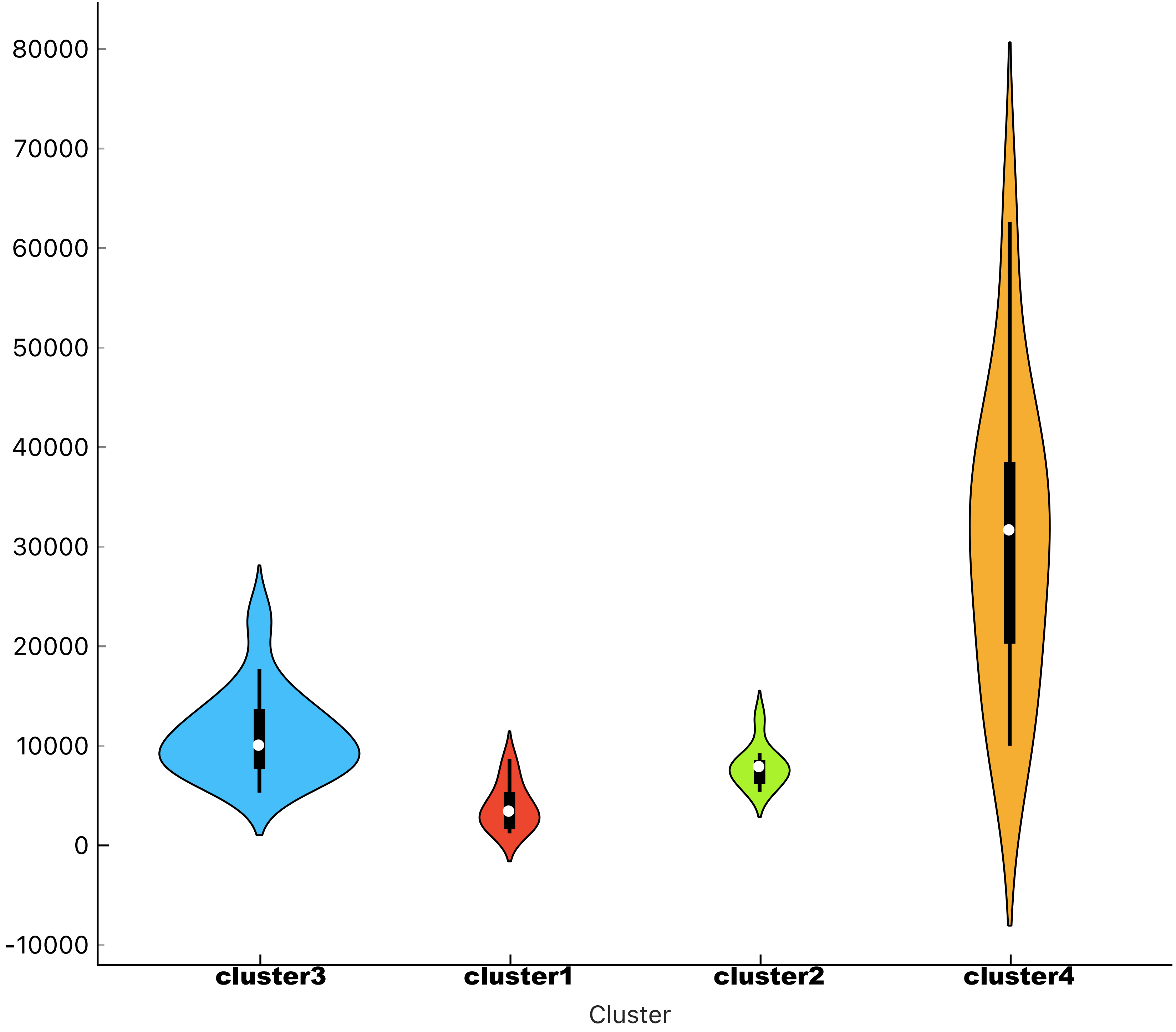

### supp info12_CBD-violin

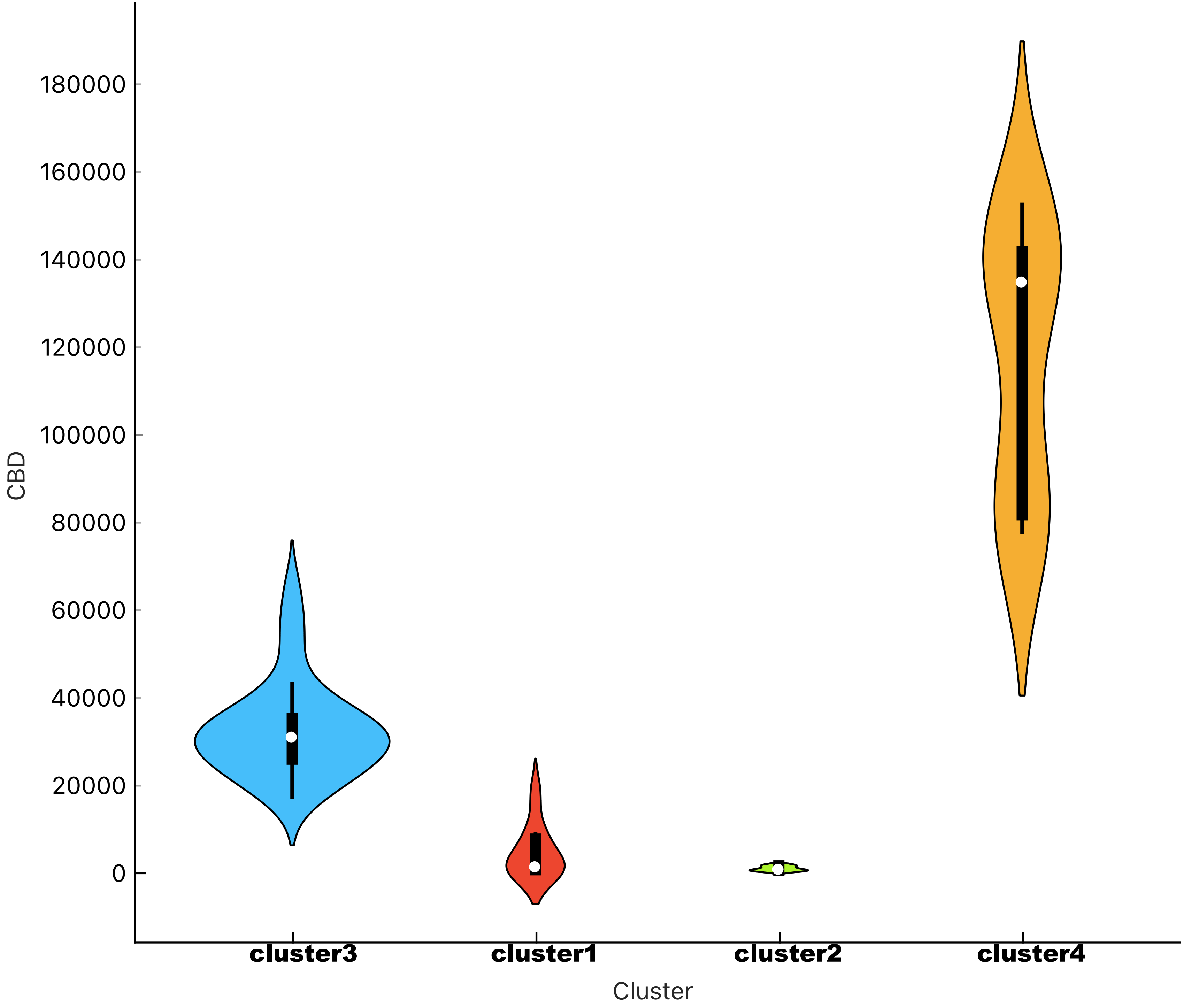

### supp info13_CBDA-violin

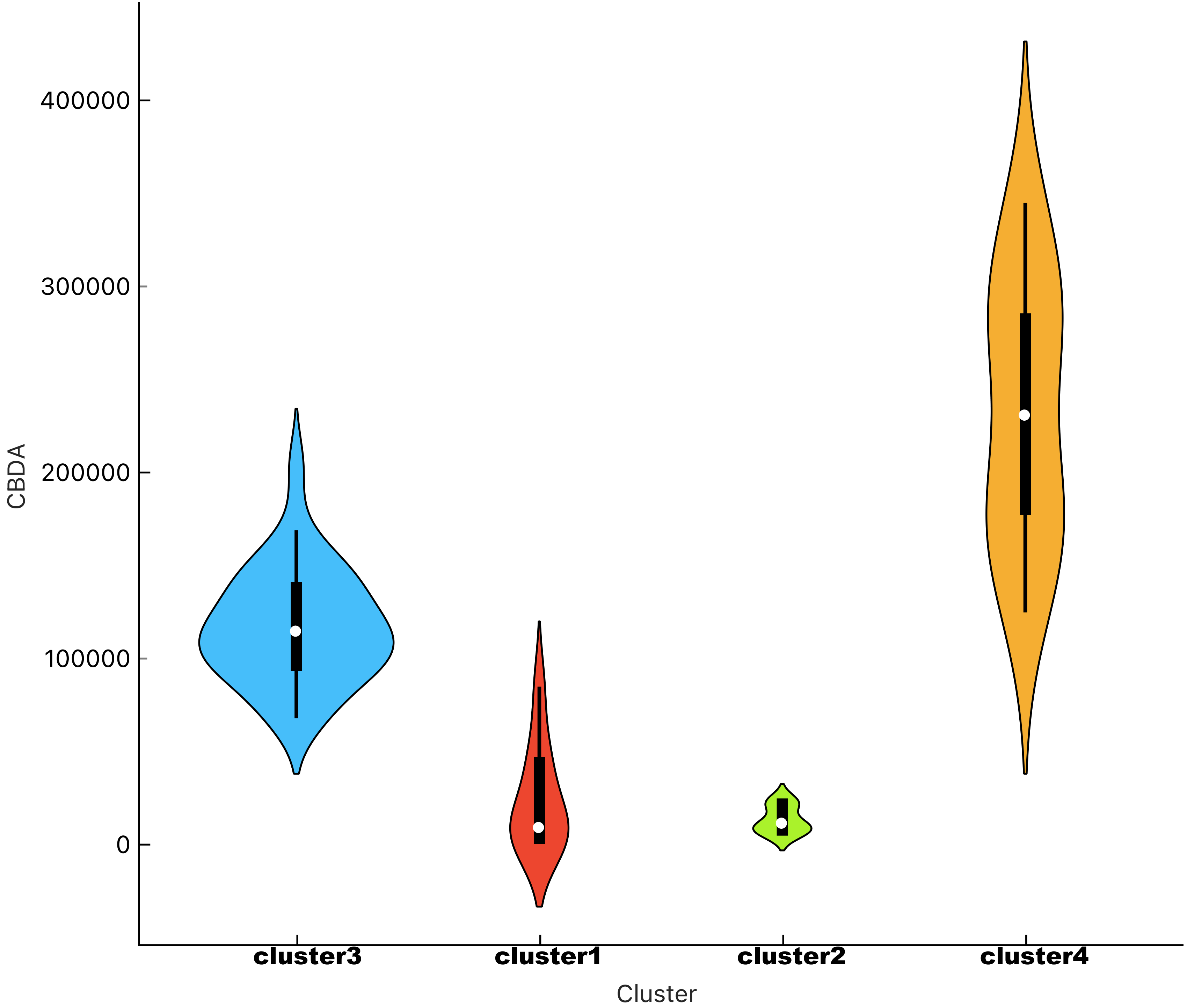

### supp info14_CBG-violin

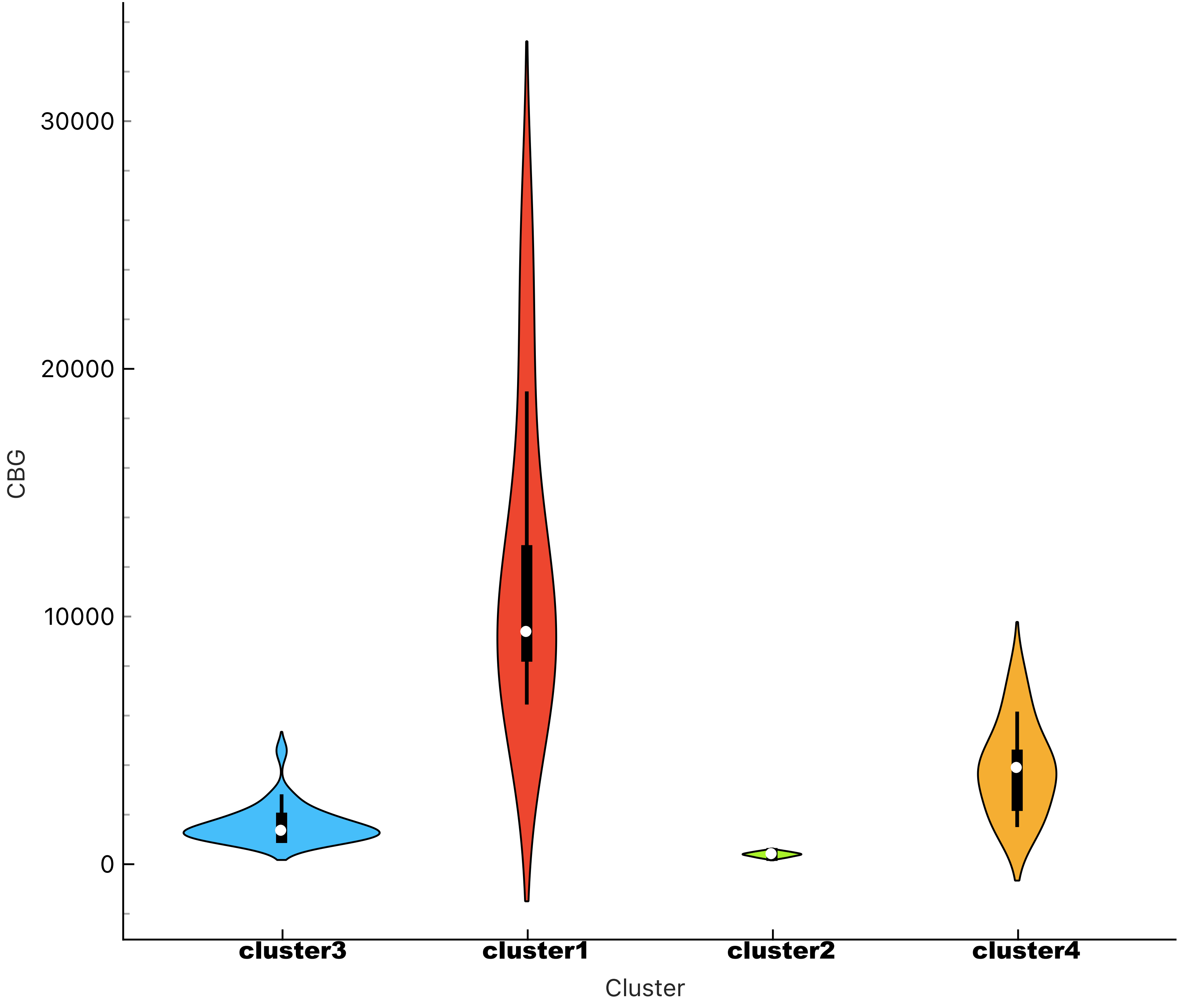

### supp info15_CBGA-violin

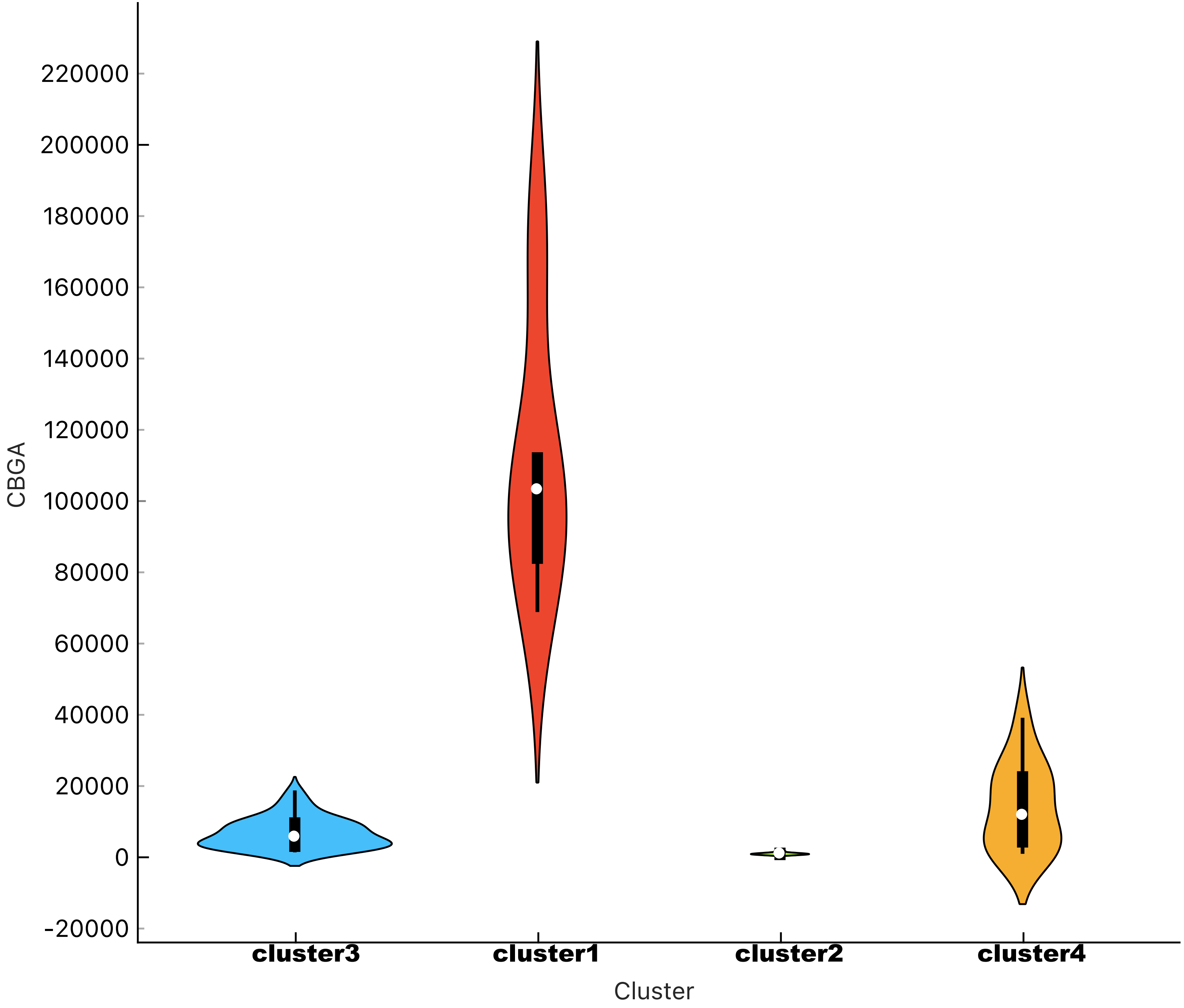

### supp info16_CBN-violin

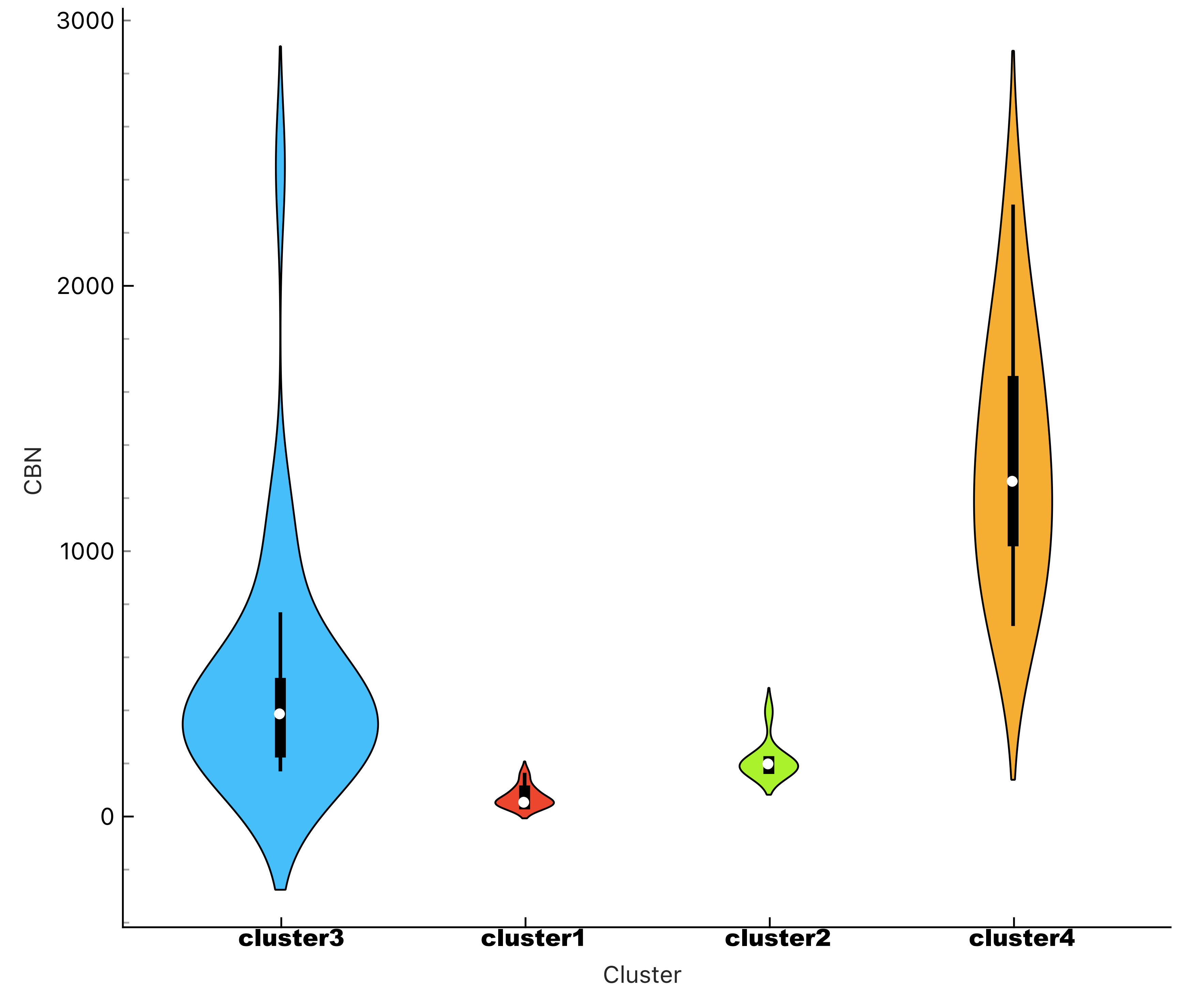

### supp info17_D8-THC-violin

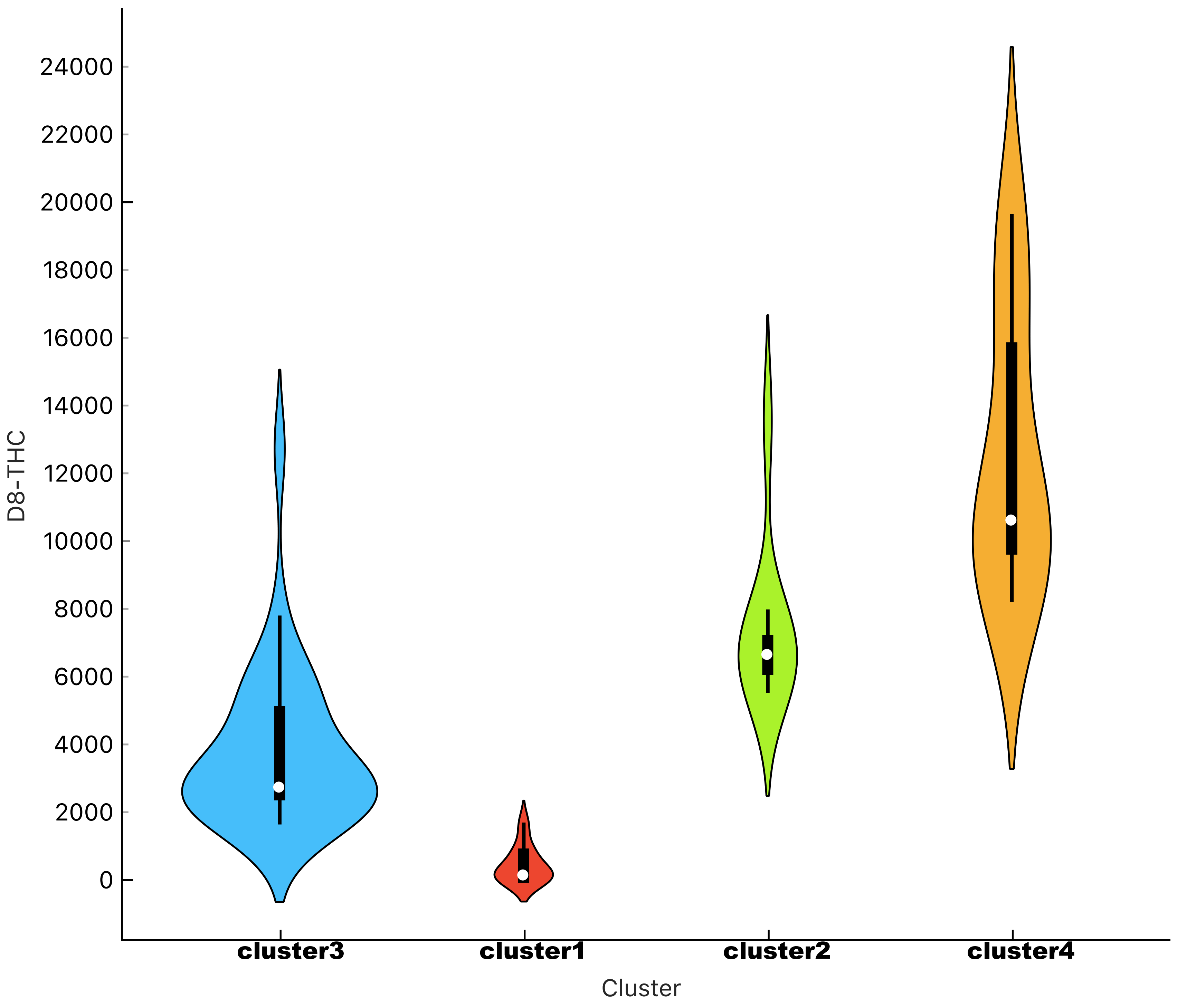

### supp info18_D9-THC-violin

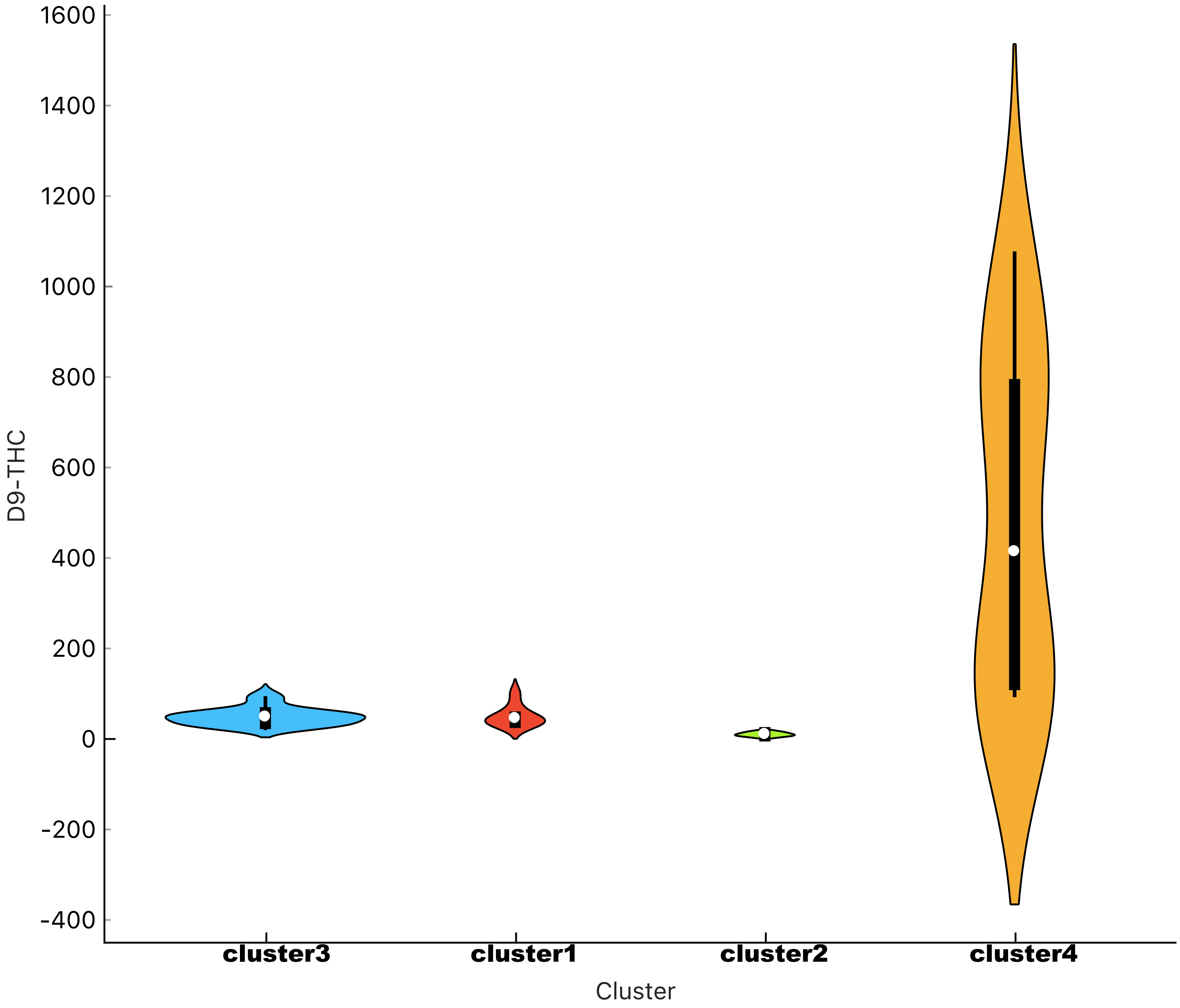

### supp info19_D9-THCA-violin

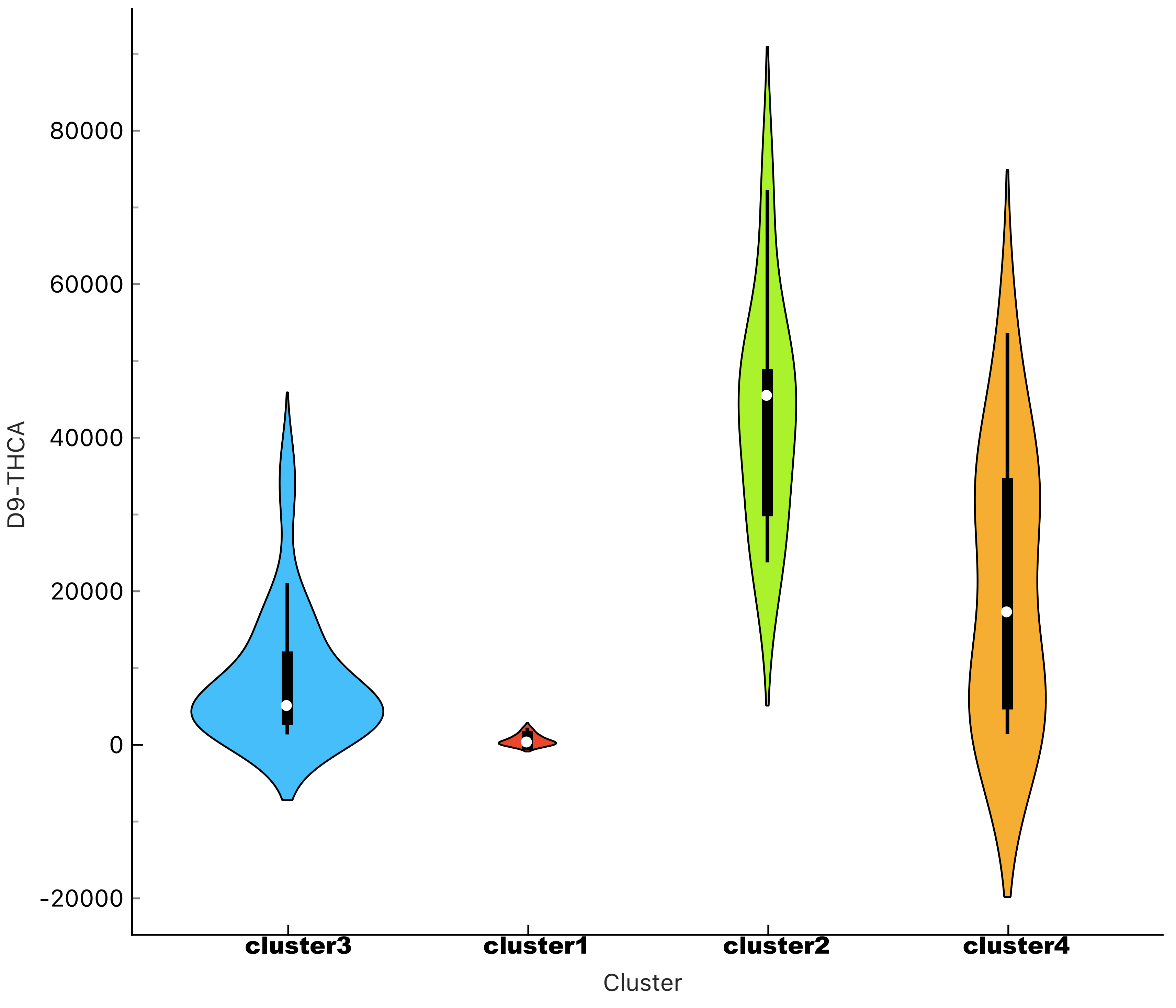

### supp info20_D9-THCV-violin

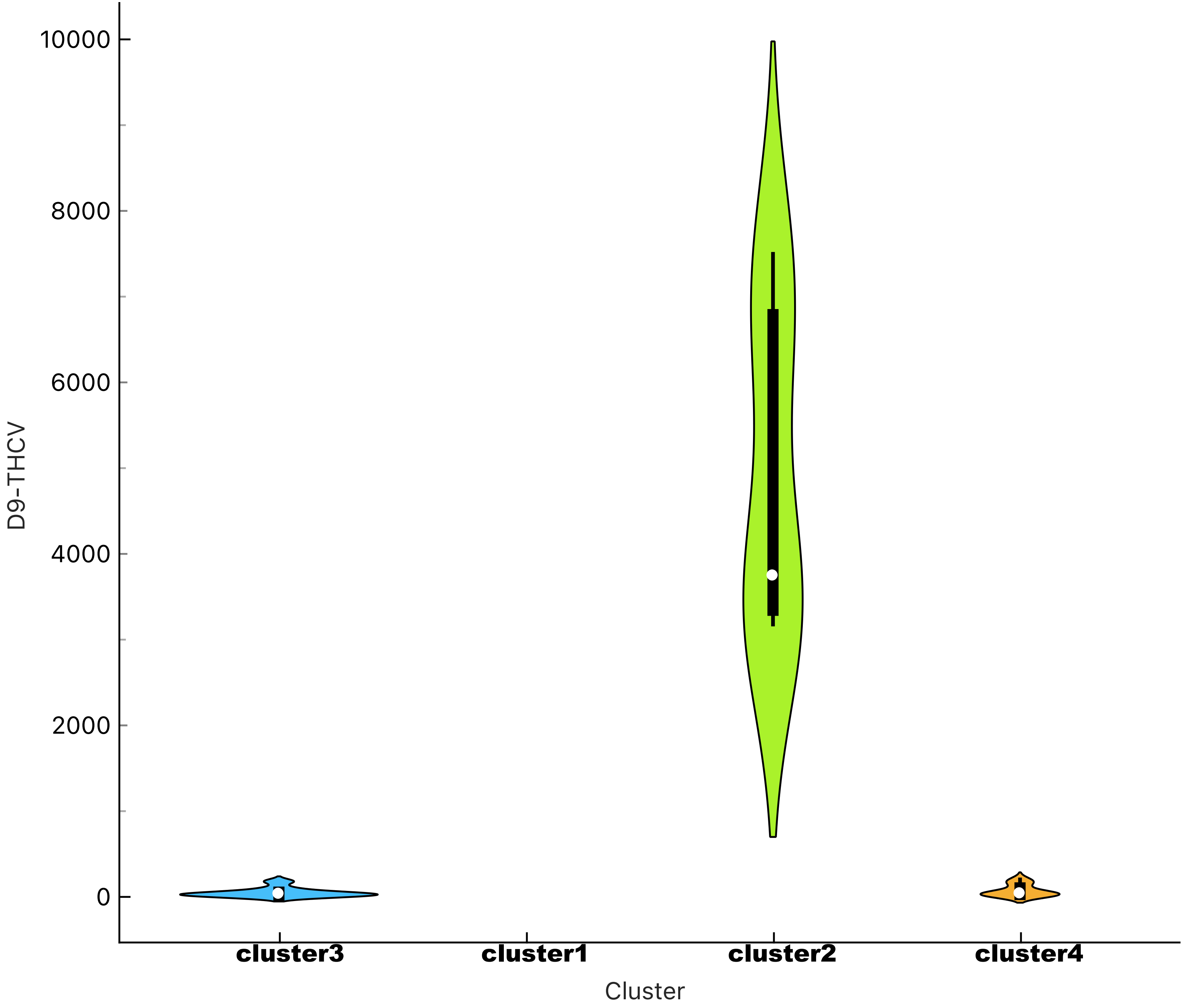
