## Supplementary material for "Cannabinoids vs. whole metabolome: relevance of cannabinomics in analyzing *Cannabis* varieties": supp info5_genotyping-structure

### Slide 1
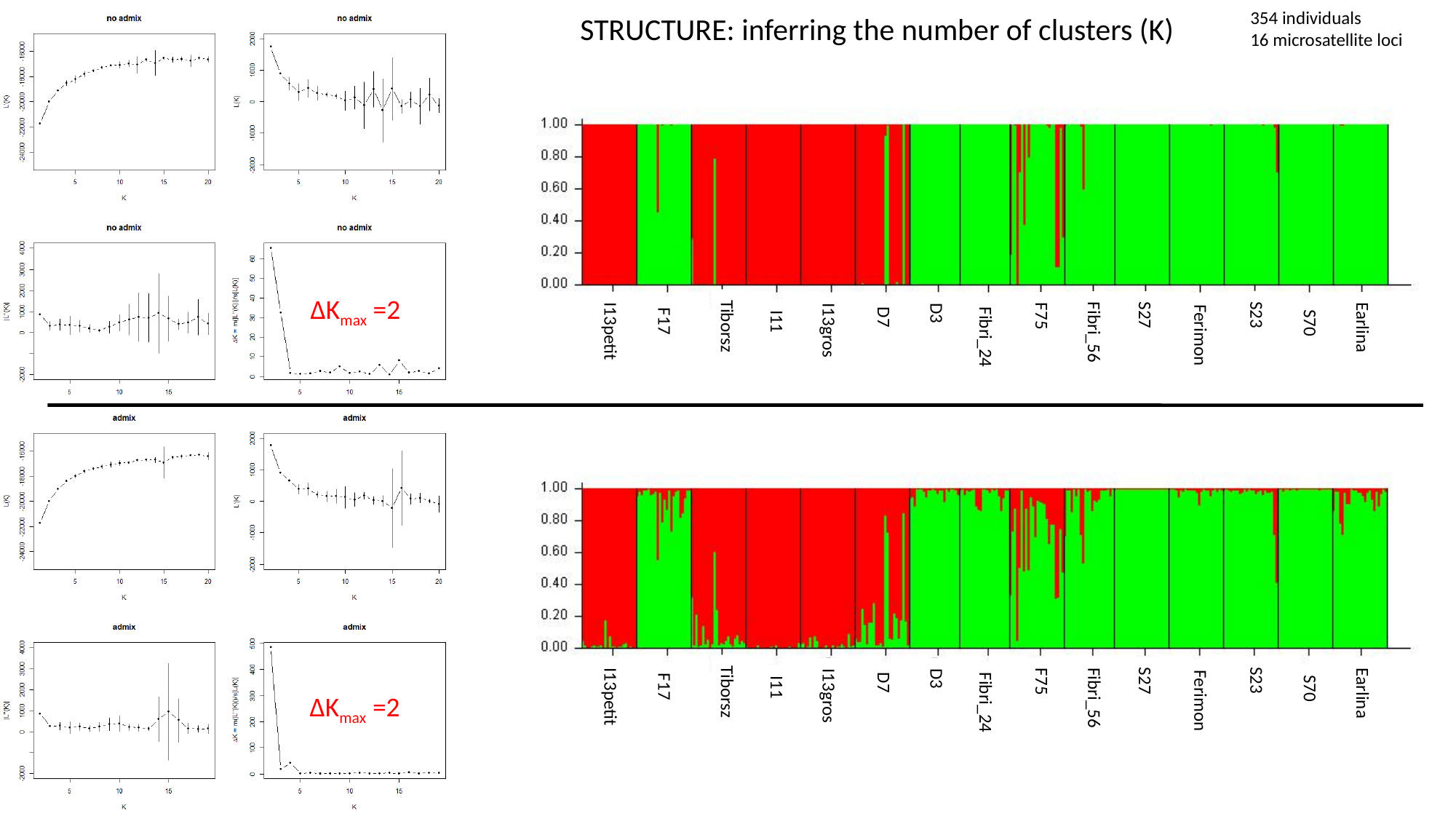

354 individuals
16 microsatellite loci
STRUCTURE: inferring the number of clusters (K)
ΔK_max_ =2
S23
S27
F75
D3
D7
S70
F17
I11
Fibri_56
Earlina
I13petit
I13gros
Ferimon
Fibri_24
Tiborsz
S23
S27
F75
D3
D7
S70
F17
I11
Fibri_56
Earlina
I13petit
I13gros
Ferimon
Fibri_24
Tiborsz
ΔK_max_ =2

### Slide 2
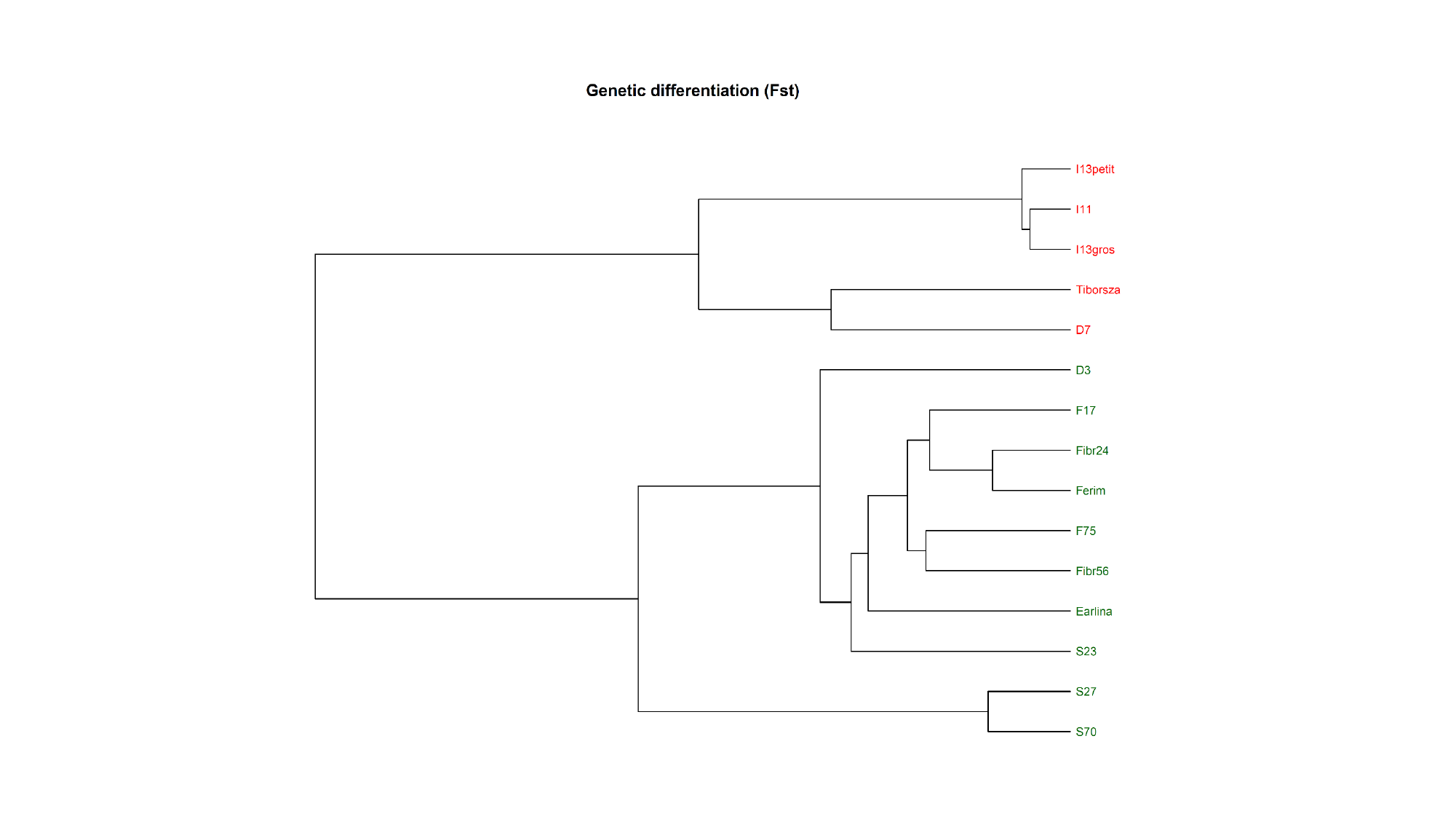
